# Prefrontal Mechanisms of Rule Learning

**DOI:** 10.64898/2026.04.01.715865

**Authors:** Rye Jaffe, Wenhao Dang, Tony Gao, Junda Zhu, Christos Constantinidis

## Abstract

Rule learning is associated with lasting changes in prefrontal activity. However, experiments typically focus on learning a single set of rules or task and there remains a significant gap regarding how related mechanisms may reflect behavioral improvements in different contexts, especially when examining across different tasks and modalities. We therefore recorded single units from chronic electrode arrays implanted in the prefrontal cortex of four monkeys as they were trained to perform spatial and object working memory tasks with the goal of assessing the resulting activity changes that would be induced by rule learning. Progression of training allowed behavioral improvements to be correlated with a variety of neural effects that could be observed across different task contexts, including increases or decreases of both firing rate and decoding, increases in the proportion of firing rate variance that was unexplained by sensory stimuli and motor actions, and the increased separation of population response trajectories in state space. Our results thus reveal how rule learning induces plasticity of prefrontal cortical activity, and the aspects of neural activity changes that were unique to individual tasks and modalities or common across them. Our results ultimately reveal new patterns of training effects, identifying the generalized prefrontal mechanisms that are responsible for rule learning.

## INTRODUCTION

Rule learning requires knowledge of the values of available alternatives and their dependence on context (Costa et al., 2015; Hamid et al., 2021; Averbeck and O’Doherty, 2022). Possible actions may then be assigned a continuously updated place within a hierarchy of values (Tomov et al., 2020; O’Doherty et al., 2021; Xu et al., 2024). Such hierarchies allow relevant actions to be selected based on whichever is deemed to have the highest values in the current state, with recent research linking these values to a variety of neural correlates in higher cortices (Averbeck and O’Doherty, 2022; Moneta et al., 2023; Grossman et al., 2026). Rules may be acquired through trial-and-error training, meaning that the prior trial’s outcome can be used to infer the best choice in the current trial (Lloyd et al., 2012). Neural correlates of rule learning have been described in the prefrontal cortex (PFC); the activity of individual neurons changes as associations and contingencies are learned (Asaad et al., 1998; Suchow et al., 2014; Jun et al., 2024). These changes may progress at variable time scales, reflecting rapid events or progressive understanding of a rule through training (Miller and Constantinidis, 2024).

Experimental studies, particularly in humans, have produced conflicting results making it difficult to identify common mechanisms of task learning. Training in paradigms that require working memory (WM) represents an illustrative case study. WM training may lead to higher levels of activity across the PFC (Garavan et al., 2000; Hempel et al., 2004), suggesting that repeated engagement with training strengthens the neural representations that underlie short-term information maintenance and manipulation. By contrast, other studies have demonstrated decreases in activity which have been interpreted as improvements in efficiency, instead (Schneiders et al., 2011; Takeuchi et al., 2013). There are also studies that have demonstrated increases or decreases in different prefrontal areas (Miller et al., 2022), and more subtle changes in neural dynamics rather than overall levels of activity, such as functional connectivity and network modularity, rather than overall changes in activity (Gordon et al., 2014; Finc et al., 2020). Neurophysiological studies in non-human primate models have not been able to resolve this debate, either, as they typically focus on one, or limited variations of a task, typically in a pair of monkeys, making it difficult to determine whether general principles exist (Sarma et al., 2016; Mansouri et al., 2020; Tang et al., 2022).

We were therefore motivated to determine the mechanisms of rule learning by examining prefrontal activity in monkeys across learning of a variety of tasks requiring spatial and object working memory. We sought to identify changes in neural activity that parallel the related behavioral improvements and determine if they were unique to the task trained or generalized across separate contexts and modalities.

## RESULTS

### Monkeys acquire sequential task elements via discretized training

Behavioral and neural data were collected from 4 monkeys (three male, one female) as they trained in one spatial and two object working memory tasks. In the spatial match-nonmatch task (Fig. 1a), the monkeys were required to maintain fixation, observe two stimuli appearing in sequence separated by delay periods, and to indicate if the two stimuli appeared at the same location or not by selecting one of two choice targets, defined by their shape (“H” or “Diamond”). The object match-nonmatch task (Fig. 1a) followed the same structure but prompted the monkey to indicate if two stimuli appearing in sequence had the same shape or not, by selecting one of two choice targets, defined by their color (“Green square” or “Blue square”). Finally, in the object choose-match task (Fig. 1a), the monkeys were required to maintain fixation and observe a single cue stimulus (either circle or triangle) that was followed by a delay period. Then a pair of choice targets (both circle and triangle) would appear, prompting the monkey to saccade to whichever matched the cue.

**Figure 1:**
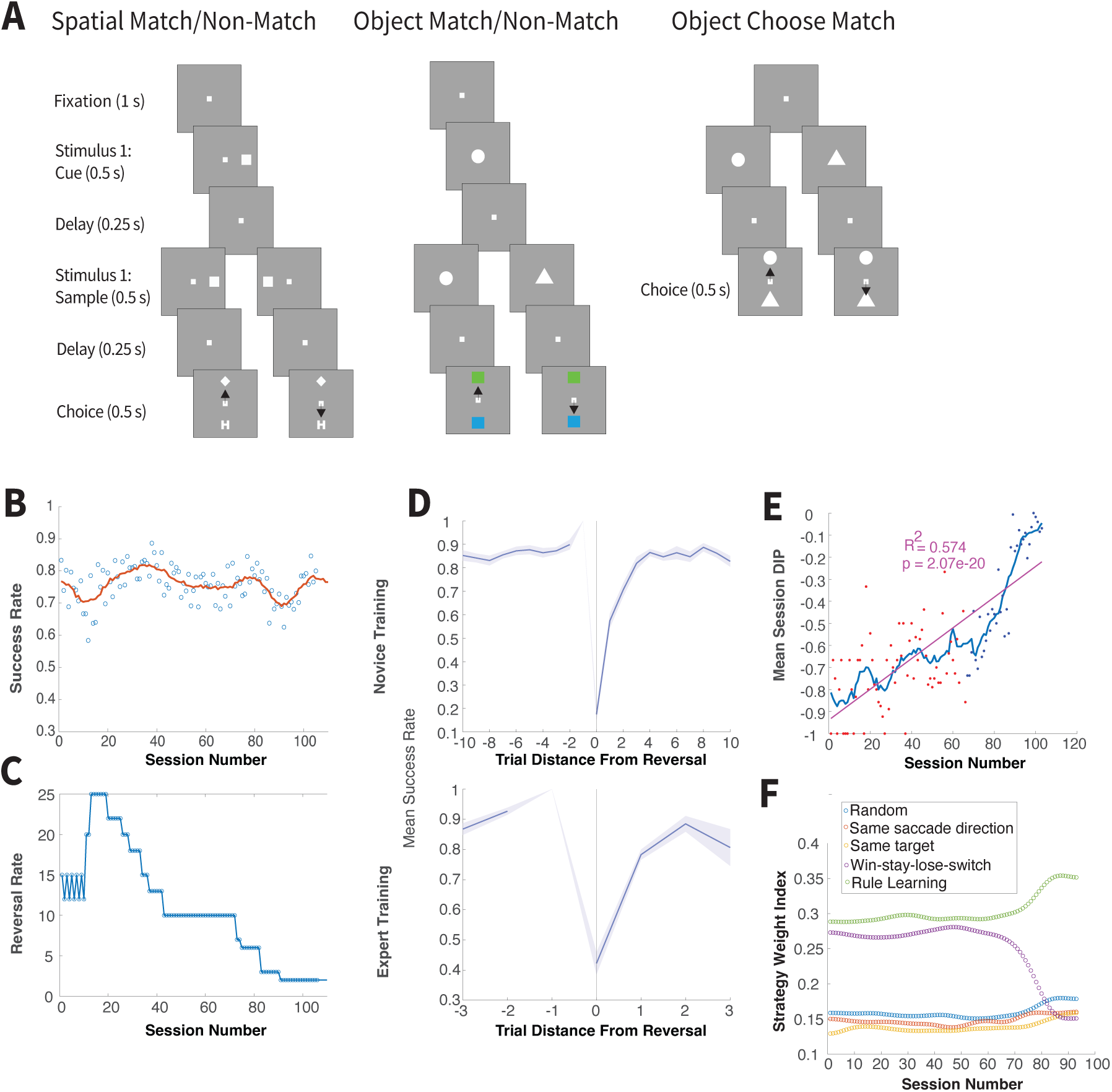
Experimental Tasks and Behavior. A. Schematic of the spatial match-nonmatch, object match-nonmatch, and object choose-match tasks. In the match-nonmatch tasks, the first stimulus (cue) remained fixed throughout training (i.e. a square on the right side of the screen for the spatial task, and a circle at the center of the screen for the object task). After a delay, this was followed by a match stimulus identical to the cue, or a nonmatch stimulus (i.e. square on the left side of the screen for the spatial task, and a triangle at the center of the screen for the object task). After a second delay, the monkey had to saccade to the “Diamond” or “Green Square” target to indicate a match or the “H” target or “Blue Square” to indicate a non-match. In the object choose-match task, the cue could be either a circle or triangle at the center of the screen. After a delay, two targets appeared (i.e. a circle and triangle) and the monkey had to saccade to whichever matched this cue. B. Mean performance of an example monkey at each session—indicated via dots—across training. Red lines indicate centered running average with a symmetric window of 5. C. Reversal rate between trial blocks for an example monkey at each session across training. Reversal rates of 2 indicate randomized reversals, by convention. D. Example monkey performance (y-axis) across trials relative to their distance from the reversal (x-axis). Shaded areas indicate standard error. The last trial that preceded the reversal was—by definition—always correct. Upper, novice; lower expert. E. Mean session DIP (y-axis) of an example monkey across training, with novice/expert training highlighted in red/blue, respectively. Thin lines represent centered running average with a symmetric window of 5. Thick lines represent linear regression of DIP on session number. F. Weight indices (y-axis) of an example monkey’s strategies across training. Each color dot represents a different strategy in one session.

Task paradigms discretized training via alternating trial blocks with different stimulus combinations. Each session of both match-nonmatch tasks was thus assigned a set number of trials that would have to be completed successfully before a reversal between trials that presented either the matching or nonmatching stimulus combination. In the case of the object choose-match task that only presented a single cue stimulus, the completion of a trial block led to a reversal between which object was used as that initial cue stimulus instead. The size of these trial blocks—also known as the “reversal rate”—was constant and manually predetermined at the start of each session. Alternating blocks of match and nonmatch trials were presented in both versions of the spatial and object match-nonmatch task, with block length decreasing over the course of multiple sessions until they were randomly interleaved, thus requiring the monkey to associate the second stimulus location/shape with the corresponding choice target to avoid failure. Alternating blocks of circle and triangle cue presenting trials were applied in each session, with block length decreasing over the course of multiple sessions until they were randomly interleaved, thus requiring the monkey to associate the cue stimulus shape with the corresponding choice target to avoid failure.

Monkeys were able to perform all three tasks with an average accuracy across sessions that was far above chance level (Fig. 1b and Fig. S2a, mean performance of spatial match-nonmatch task: 70.30%, mean performance of object match-nonmatch task: 67.96%, mean performance of object choose-match task: 66.50%) even as we progressively reduced the reversal rate over the course of training. This indicates that all monkeys were fully participating in the task and progressively acquired the rule needed to perform the randomly interleaved task for both spatial and object information.

During the first training sessions when the monkey had not yet begun to learn the new rules of this experiment, they were expected to perseverate and would thus almost always fail the trial in which there was a reversal between matching and nonmatching stimuli. This led to a consistent Drop In Performance (henceforth abbreviated as DIP) at the reversal trial (Fig. 1d and Fig. S2c). This forced the monkeys to realize that a new element had been added to their task: the rewarded target could now change over the course of a session. The magnitude of DIP was calculated for each session by subtracting from the mean success rate of the trials where the reversal occurred the mean success rate of trials that occurred directly prior to this reversal.

Monkeys were given no indication for when a reversal occurred, and so over the course of additional training sessions, DIP would only decrease in absolute magnitude if they began to learn that when trials changed from presenting pairs of matching stimuli to presenting pairs of nonmatching stimuli (or the reverse), the rewarded choice target would be change as well, switching from the Diamond and Green targets to the H-shape and Blue targets (or the reverse). Learning this rule allowed the monkeys to adjust their behavior accordingly to avoid failure. As a result, the magnitude of DIP could be used to identify the stage of learning. We relied on DIP to classify sessions into “novice” and “expert” around the point where the DIP running average began the longest path of monotonic increase (Fig. 1e and Fig S2d).

During training, the reversal rate was gradually reduced across multiple sessions, shortening trial blocks over the course of training until the presentation of match or nonmatch stimulus (i.e. trials presenting stimulus pairs with the same location/shape. in the spatial/object match-nonmatch tasks, or trials presenting the circle and triangle stimulus in the object choose-match task), were randomly interleaved. A complete lack of understanding of the rule would result in failure following a reversal between the presentation of match or nonmatch stimulus, thus causing a consistent drop in performance (DIP). In contrast, full understanding of the rule would result in no DIP when the rewarded choice target reversed. We thus tested for evidence of rule learning as training progressed by determining whether the absolute DIP magnitude decreased as a function of training. This was indeed the case. An example is shown in Fig. 1e (linear regression; F(1,101) = 135.9, p=2.07E-20)), The rest of the cases are shown in Fig S2d (p<E-4 in all cases).

Our training paradigm was designed to incentivize genuine rule learning for success and so our monkeys were forced to use trial-and-error to realize that the stimuli presented prior to the choice targets would determine which of these targets would be rewarded. In behavioral terms, this meant that the optimal strategy—that is, the only strategy that could potentially achieve a consistent 100% success rate—would be to examine the presented stimuli, and then saccade to the associated choice target. However, at least during the start of training, a variety of alternative task strategies could also be used to produce reward, even in the absence of rule learning. For example, longer trial blocks preceding a reversal between the presentation of match or nonmatch stimulus (i.e. trials presenting stimulus pairs with the same location/shape. in the spatial/object match-nonmatch tasks, or trials presenting the circle and triangle stimulus in the object choose-match task) would allow the monkey to continue receiving reward by saccading to the same reward target for a longer span of time, without needing to learn the difference between trials with different stimulus presentations. Once these trial blocks came to an end, the monkey could then recognize the sudden halt in reward as a signal to shift their saccade to the other choice target instead, thus allowing them to continue receiving reward without genuine rule learning. This was known as the “Win-stay-lose-shift” strategy.

Other strategies that could serve as possible alternatives to genuine rule learning included choosing the target at random, regular alternation between targets with each trial, choosing to saccade towards the same location every trial regardless of which target was at that location, or choosing the same target stimulus every trial regardless of the target’s location—just as the monkeys had learned to do during the pre-training paradigm.

We assessed the weight index (see Methods) for each alternative strategy across each session by comparing the percentage of trials in which the monkey’s choice was the same as the choice that would be dictated under each of the given strategies to determine the probability of each strategy being used, relative to the others (Fig. 1f and Fig. S2e). As expected, the weight index of the optimal rule learning-based strategy increased significantly from novice to expert training to far surpass all other strategies by the end of training (two-way ANOVA, F(4,515)=85.95 for interaction of novice/expert training and strategy type; p=6.8E-56 in Fig. 1f; p<E-4 for all other cases shown in Fig. S2e).

### Firing rate changes during training

Extracellular neurophysiological recordings were recorded from the lateral PFC of the four monkeys with chronic arrays (Fig. S3A-D). A total of 270 single units were recorded over the course of 159 sessions from two monkeys assigned to the spatial match-nonmatch task, 1348 single units were recorded over the course of 203 sessions from one monkey assigned to the object match-nonmatch task, and 1765 single units were recorded over the course of 127 sessions from two monkeys assigned to the object choose-match task (a total of 3383 single units and 489 sessions across all tasks).

We sought to examine whether rule learning could be characterized by changes in firing rate across all tasks. Previous neurophysiological studies have suggested an overall increase in mean firing rate as learning progresses (Tang et al., 2022). However, such studies have also revealed decreases in firing rate during acquisition of some task elements (Tang et al., 2019; Tang et al., 2022). For this reason, we quantified prefrontal firing rates in each epoch of each task to assess how activity changed as training progressed. Mean firing rates in correct trials across all neurons were plotted (Fig. S3E). We then examined the changes in mean firing rate between the “novice” and “expert” parts of training, separately for each task, epoch, and monkey (Fig. S3F). We used the raw firing rate for the fixation epoch and changes relative to this baseline fixation rate for the rest of the epochs by subtracting the baseline firing rate from the firing rate of other epochs (He, 2013). This analysis revealed no consistent changes in firing rate with training. The choice epoch had the strongest linear relationship across all monkeys/tasks, showing an overall decrease in firing after training, though even in this case the relationship was weak (Pearson correlation coefficient r=0.106). Firing rate changes across tasks and monkeys in other epochs exhibited an even weaker or negligible linear relationship (fixation/cue/first delay’s Pearson correlation coefficient r = 0.010, 0.031, 0.030).

We refined our analysis by examining the possibility that a systematic relationship in firing rate changes between tasks might be non-linear and might progress as a function of training. We therefore relied on a Generalized Additive Mixed Models approach, using DIP as the dependent variable in this analysis (Fig. 2). This analysis serves as a flexible, semiparametric method for identifying and estimating nonlinear effects of covariates on the outcome variable (i.e. firing rate), while accounting for the possibility that differing firing rate intercepts from different monkeys might obscure the relationship that we were seeking to examine. We first examined the baseline fixation epoch firing rate across all available cells, revealing a slight trend of increase (GAMM, baseline fixation: F = 0.078, p = 0.045). Repeating this analysis in other task epochs, after subtracting the baseline firing rate, revealed a highly significant nonlinear trend of decrease in firing rate of the choice epoch (GAMM: p = 0.001, F = 3.62). Firing rate changes in other epochs were not significant (GAMM, cue: p = 0.776, first delay: p = 0.696).

**Figure 2:**
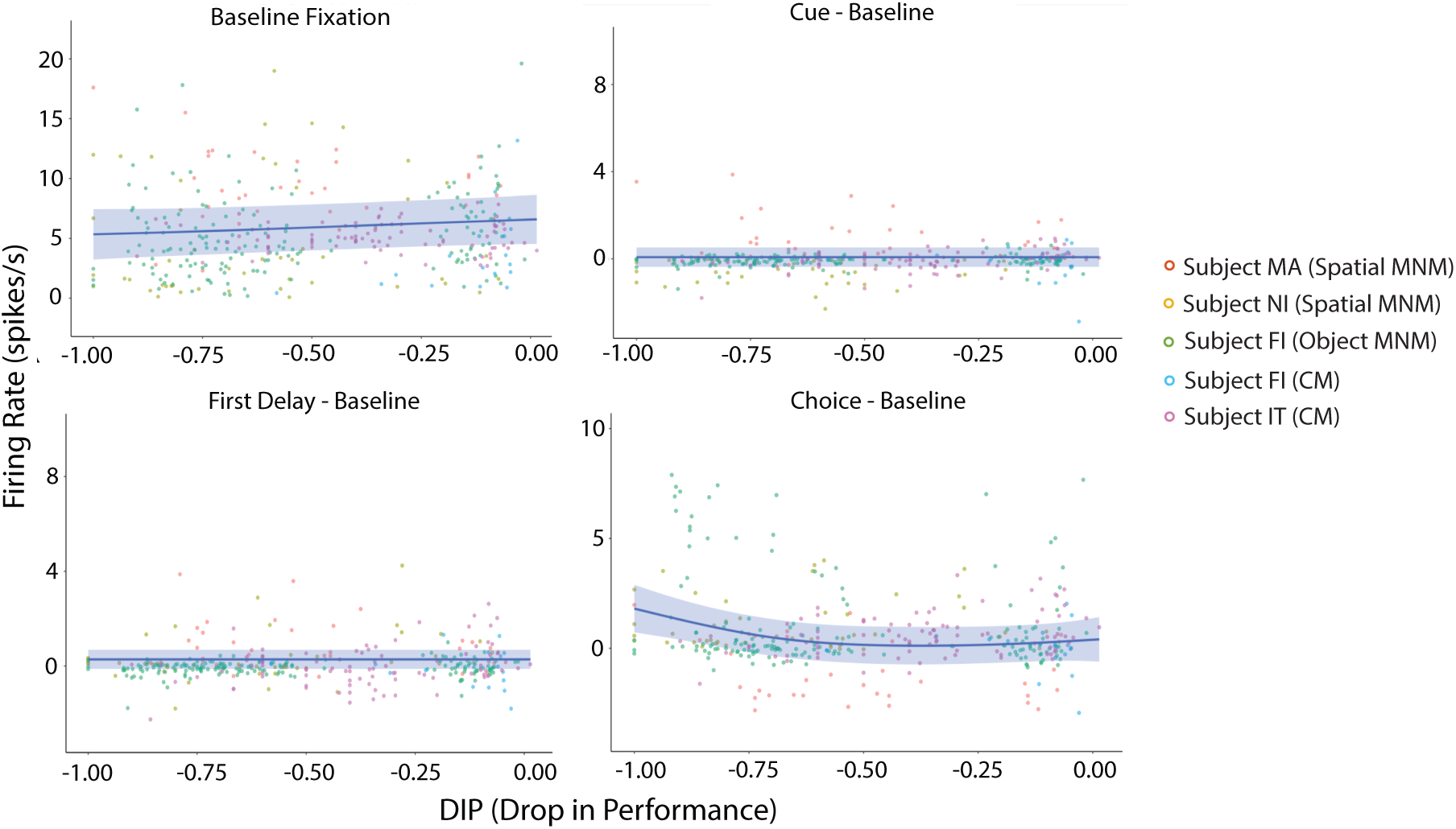
Nonlinear training effects on firing rate. The trajectory of the mean session firing rate (y-axis) in different epochs (baseline fixation, as well as cue epoch, first delay epoch, and choice epoch after subtracting the mean baseline fixation activity) across mean session DIP (x-axis). Solid lines indicate training effects of the GAMM fittings. Shaded ribbons denote the 95% confidence intervals (CIs). Each dot represents a single session, and individual monkeys are indicated by different colors. Outliers were removed from the visual plots by scaling the y-axis, with a total of 2 outliers removed from the fixation epoch plot, 1 outlier removed from the cue epoch plot, 1 outlier removed from the first delay epoch plot, and 2 outliers removed from the choice epoch plot for better visualization, though these points were still used in the curve estimation.

These results suggest that the overall increase in firing rate after training reported in previous studies is not a general phenomenon across different tasks, with only a minor increase in baseline firing rate captured by the GAMM across our experiments. Instead, a decrease in firing rate during the choice epoch period appeared to have a more consistent effect.

### Training increases unexplained variance of firing rate

In addition to examining changes in overall firing rate, we sought to examine the hypothesis that rule learning would be characterized by changes in the variance of firing rate explained by task variables, as learning takes place. Monkeys trained in the two-armed bandit reversal learning task would display increased unexplained variance in firing rate during the change between conditions in a reversal learning task (Bartolo and Averbeck, 2020). This means that when firing rate was modeled with known task variables, the amount of variance that could not be explained by this model would significantly increase during the trials when the monkeys made an internal decision to reverse their target preference. We therefore sought to determine whether this increased unexplained variance would occur over the process of rule learning as well, or alternatively, if this might be a phenomenon that would only emerge after the completion of training.

We therefore used a linear regression to model firing rate as a combination of three task factors a) the presentation of match or nonmatch stimulus (i.e. trials presenting stimulus pairs with the same location/shape in the spatial/object match-nonmatch tasks, or trials presenting the circle and triangle stimulus in the object choose-match task); b) the direction of the rewarded choice target (i.e. whether the rewarded target appeared up or down), and c) whether the trial was answered correctly or not (Fig. 3; see also the insets of Fig. 4 for an illustration of the match/nonmatch, and target direction conditions). Firing rate was thus expressed as:

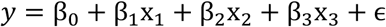

where *y* is firing rate calculated in a 400 ms bin, and *x_1_, x_2_,* and *x_3_* are categorical variables representing the predictors of matching or nonmatching status, up or down direction of the correct target, and correct or error outcome of the trial. The amount of firing rate variance that this model failed to explain across all trials in a session was calculated as 1-R^2^. Such variance would be implied to come from firing rate variability that is not linearly related to the three variables of our model. Values computed in successive 400 ms bins tiling the choice epoch (and the entire trial) were then averaged to calculate the values depicted in Fig. 3 (and Fig. S4).

**Figure 3:**
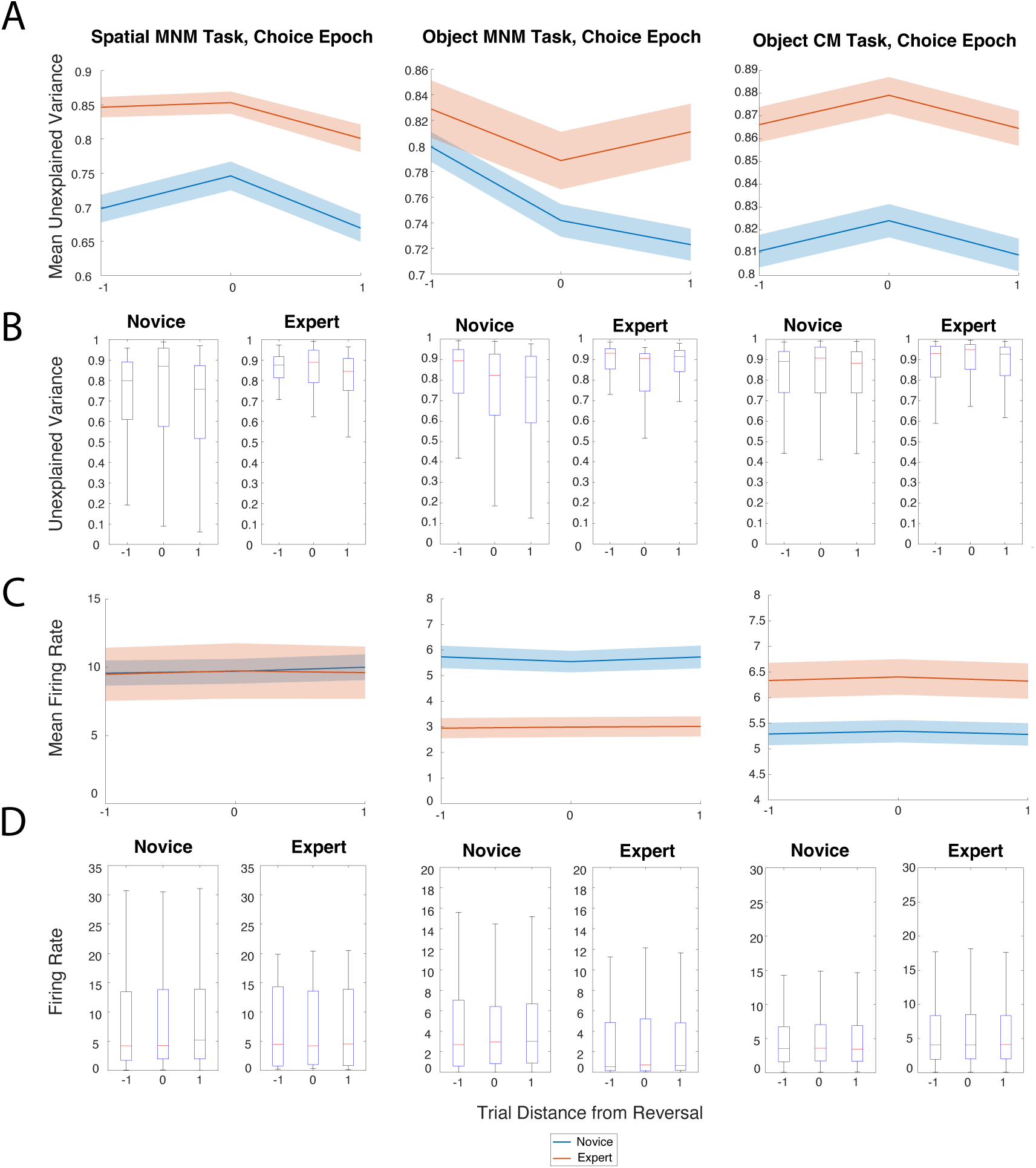
Training increases unexplained variance across tasks. A. Mean unexplained variance of firing rate in the choice epoch across trials relative to their distance from the reversal. Shaded areas indicate standard error. Blue indicates novice while red indicates expert stage. B. The same data as in A represented as boxplots. On each box, the central mark indicates the median, and the bottom and top edges indicate the 25th and 75th percentiles, respectively. The whiskers extend to the most extreme data points not considered outliers—defined as any values that were more than 1.5 * the interquartile range (i.e. the distance between the 25th and 75th percentiles) below the 25^th^ percentile, or above the 75^th^ percentile. C-D. Same conventions as A-B but examining choice epoch firing rate instead of unexplained variance.

**Figure 4:**
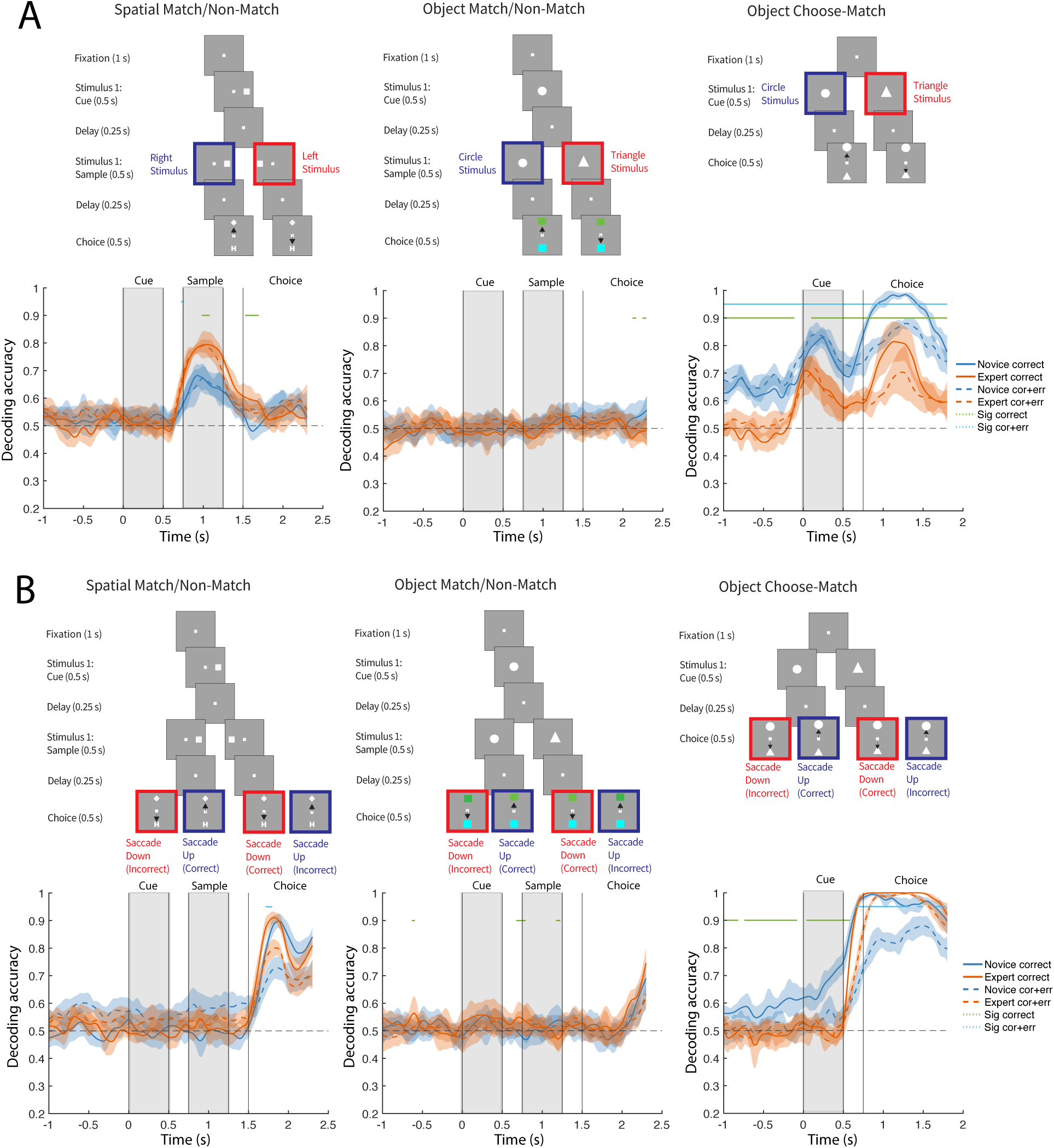
Training has no systematic pattern in decoding effects across tasks. Top diagrams illustrate A. the presentation of match or nonmatch stimulus (i.e. trials presenting stimulus pairs with the same location/shape in the spatial/object match-nonmatch tasks, or trials presenting the circle and triangle stimulus in the object choose-match task), and B. the monkey’s chosen saccade direction (up or down) for each task. Bottom plots illustrate the accuracy of decoding this task factor from the novice (blue) versus expert (red) neurons over the time course (x-axis) of each task. Vertical black lines indicate each task epoch. Solid blue and red lines indicate the use of exclusively correct trials for training and testing data. Dashed blue and red lines indicate the use of both success and error trials for training and testing data. Shaded ribbons represent 95% confidence intervals. Significant differences between novice and expert decoding are marked overhead in green and cyan (permutation test, p < 0.04). We performed our decoding analyses using an equal number of cells drawn from novice and expert training sessions (Fig. S6). This was a total of 100 novice and expert cells from the spatial match-nonmatch task (left), 400 novice and expert cells from the object match-nonmatch task (middle), and 400 novice and expert cells from the object choose-match task (right).

Unexplained variance was examined across a range of 3 trials around the reversal (i.e. the trial directly before, during, and after the reversal), and so cells required a minimum of three trials at each position to be included in this analysis. We then used a two-way ANOVA to compare the response vector of this unexplained variance across the grouping vectors (factors) of training level (i.e. novice vs expert), and trial distance relative to the reversal. We focused particularly in the choice epoch, after all the stimulus information had been revealed and the subject received feedback on its choice, in the form of reward, or absence thereof (Fig. 3). This analysis revealed a significant main effect for training stage where unexplained variance was observed to increase from novice to expert training (spatial match-nonmatch task: F(1, 600) = 37.29, p<E-4; object match-nonmatch: F(1, 1365) = 14.6, p = 0.0001; object choose-match: F(1, 2937) = 71.3, p<E-4). Unexplained variance also tended to be higher for the trial where the reversal occurred, however the difference between trials arranged relative to the reversal was not consistent across tasks (spatial match-nonmatch task: F(2, 600) = 3.11, p = 0.045; object match-nonmatch: F(2, 1365) = 4.98, p = 0.007; object choose-match: F(2, 2937) = 2.04, p = 0.131). There was no significant interaction between early/late training and serial position in either of the three tasks (spatial match-nonmatch: F(2, 600) = 0.32, p = 0.726; object match-nonmatch: F(2, 1365) = 1.47, p = 0.231; object choose-match: F(2, 2937) <E-4, p = 0.999).

Repeating the same two-way ANOVA for unexplained variance averaged across time bins spanning all epochs showed similar results (Fig. S4). A significant main effect for training stage was observed, where unexplained variance increased from novice to expert training (spatial match-nonmatch: F(1, 600) = 42.61, p<E-4; object match-nonmatch: F(1, 1365) = 30.28, p<E-4; object choose-match: F(1, 2937) = 95.61, p<E-4).

This pattern of unexplained variance increase with training was not tied to systematic firing rate differences between training stages. We examined choice epoch firing rate across the same range of 3 trials around the reversal (Fig. 3C-D), using a similar two-way ANOVA to compare the response vector of firing rate across the grouping vectors (factors) of training level (i.e. novice vs expert), and trial distance relative to the reversal. This analysis revealed no consistent effects across tasks. There was no significant difference in firing rate for the trials examined in the spatial match-nonmatch task (F(1, 600) = 0.02, p = 0.893). Lower firing rate was observed for the expert stage in the object match-nonmatch: F(1, 1365) = 35.8, p<E-4. Higher firing rate was observed the expert stage in the object choose-match: F(1, 2937) = 20.53, p<E-4. No significant effect of trial distance from the reversal was observed for any task, either (spatial match-nonmatch: F(2, 600) = 0.02, p = 0.979; object match-nonmatch: F(1, 1365) = 0.02, p = 0.982; object choose-match: F(1, 2937) = 0.04, p = 0.963). Repeating this analysis with firing rate averaged across all epochs yielded similar results as well (Fig. S4C-D). These findings suggest that the consistent increase in unexplained variance we observed after training was not dependent on systematic firing rate changes across tasks/monkeys.

We further wished to test which of the three task variables that we used in our regression model changed the most with training. To evaluate this, we calculated partial R square for the three task variables; The explanatory power of reward status (i.e. whether the trial was rewarded or unrewarded) diminished markedly as training progressed and the animal internalized the task rules, leading to more predictable outcomes. This was true across all task paradigms, while the contributions of the remaining variables remained stable (Fig. S5). These results suggest that reward status account for a greater proportion of firing rate variance early in the training process, when the subjects determine their responses based on these factors, on a trial-by-trial basis. These factors become less important once the monkey understands the task rule, and the appropriate response becomes deterministic, and particularly the reward outcome does not inform the choice to be selected in the next trial.

### Decoding analysis

To understand how training may affect the various types of task-related information that can be represented in prefrontal populations, we performed a decoding analysis. Decoding accuracy has been previously shown to increase for some specific task factors over the course of training, such as trials presenting matching versus nonmatching stimuli, though importantly, decoding has also been shown to remain constant for other specific task factors, such as the shape or location of these presented stimuli (Meyers et al., 2012).

We first sought to decode the match or nonmatch status of the sample stimulus. We performed this analysis both based on correct trials alone, and including correct and error trials. Decoding performance was compared between novice and expert training using a nonparametric randomization (permutation) test. As was the case with linear changes in firing rate, we did not observe a systematic pattern of decoding changes across tasks as rule learning took place over the progression from novice to expert training. Rather, differentiating between a match and a nonmatch stimulus appeared to involve three different decoding patterns. In the spatial match-nonmatch task, we observed that the progression from novice to expert training led to significant (permutation test: p < 0.04) improvements in decoding the presentation of match or nonmatch stimulus (Fig. 4A-left)). This improvement was most evident when the classifier was trained/tested with exclusively correct trials (solid lines in Fig. 4) than when error trials were also included (dotted lines in Fig. 4). In other words, neural activity after training better differentiated whether the second stimulus was presented on the right or left during the second stimulus presentation epoch. In contrast, no such differentiation was clearly evident in the object match-nonmatch task (Fig. 4A-center), regardless of whether error trials were included or not. Across our sample of neurons from this task, very few conveyed information about whether the second stimulus was a circle or triangle, both early and late in the training process. Finally, the object-choose-match task was characterized by the appearance of a bias in neural activity, early, which disappeared as training progressed (Fig. 4A-right). This meant that neurons represented whether the monkey would indicate the circle or triangle to be the match even before the sample was presented (as it was possible to do, since these trials were presented in blocks with the same cue stimulus). This decoding bias occurred across the full span of the trial, regardless of whether error trials were included or not. Similar patterns were evident for the decoding of the monkey’s choice to saccade up or down. In the spatial match-nonmatch tasks, we observed that the progression from novice to expert training led to significant (permutation test: p < 0.04) improvements in decoding the chosen direction of the monkey’s saccade (Fig. 4B-left). Decoding accuracy was greater in correct trials, than when both correct and error trials were used to train and test the classifier. By contrast, no improvement in decoding accuracy of the saccade direction was clearly evident in the object match-nonmatch task, regardless of whether error trials were included or not (Fig. 4B-center). In the choose-match task, a bias was present again (Fig. 4b-right). Neurons represented early whether the monkey would choose to saccade up or down even before the targets were presented. This bias was most evident when the classifier was trained/tested with exclusively correct trials.

Decoding of other variables such as the configuration of the choice targets (i.e. whether a specific choice target appeared up or down regardless of whether this was the rewarded target); the monkey’s choice to saccade for a match (regardless of whether this choice was correct); the direction of the rewarded choice target (regardless of whether the monkey chose to saccade to this target); and the combination of a match or nonmatch stimulus presentation with the direction of the rewarded target are depicted in Fig. S6.

To summarize, our results indicate that several different patterns of neuronal activity changes supported learning of different tasks. In the spatial match-nonmatch task, they relied upon representing the match or nonmatch stimulus, which gradually improved over the course of training. In the shape match-nonmatch task, the relative lack of decoding is less clear. Prior studies that observed relatively few cells representing object stimuli in the PFC may suggest this lack of decoding to be due to a so-called “needle-in-the-haystack” problem wherein the neural representations of the match or nonmatch stimulus may rely upon a small number of highly selective neurons that our sampling methods seemed to be largely unable to gather (Dang et al., 2022). Finally, in the object choose-match task, the pre-training paradigm was suggested to have led our monkeys to develop a strong bias towards a single saccade direction—even prior to the stimulus presentation—which gradually faded over the course of training as reversal rates were shortened and monkeys lost the ability to predict trials in advance.

### Principal component analysis

While the decoding analysis discussed in the preceding section was a supervised analysis that identified relationships between neural activity and known task variables, unsupervised methods such as principal component analysis (PCA) can reveal training effects on latent population dynamics and neural trajectories that might not have been evident from our decoder analysis alone (Mante et al., 2013; Mozumder and Constantinidis, 2023). We therefore performed a targeted PCA analysis (Mante et al., 2013), which decomposed the matrix of neural activity along the axis coding the match or nonmatch status (or shape of the cue in the choose match task), as well as along the axis coding the direction of the rewarded choice. The average population response for a given condition and time was represented as a point in state space projected into the two-dimensional subspace (Fig. 5A-B). Full neural trajectories from the empirical novice and expert training data are shown in Fig. S7. We then compared trajectories in the state space defined by these components across novice and expert training (Fig. 5 and Fig. S7).

**Figure 5:**
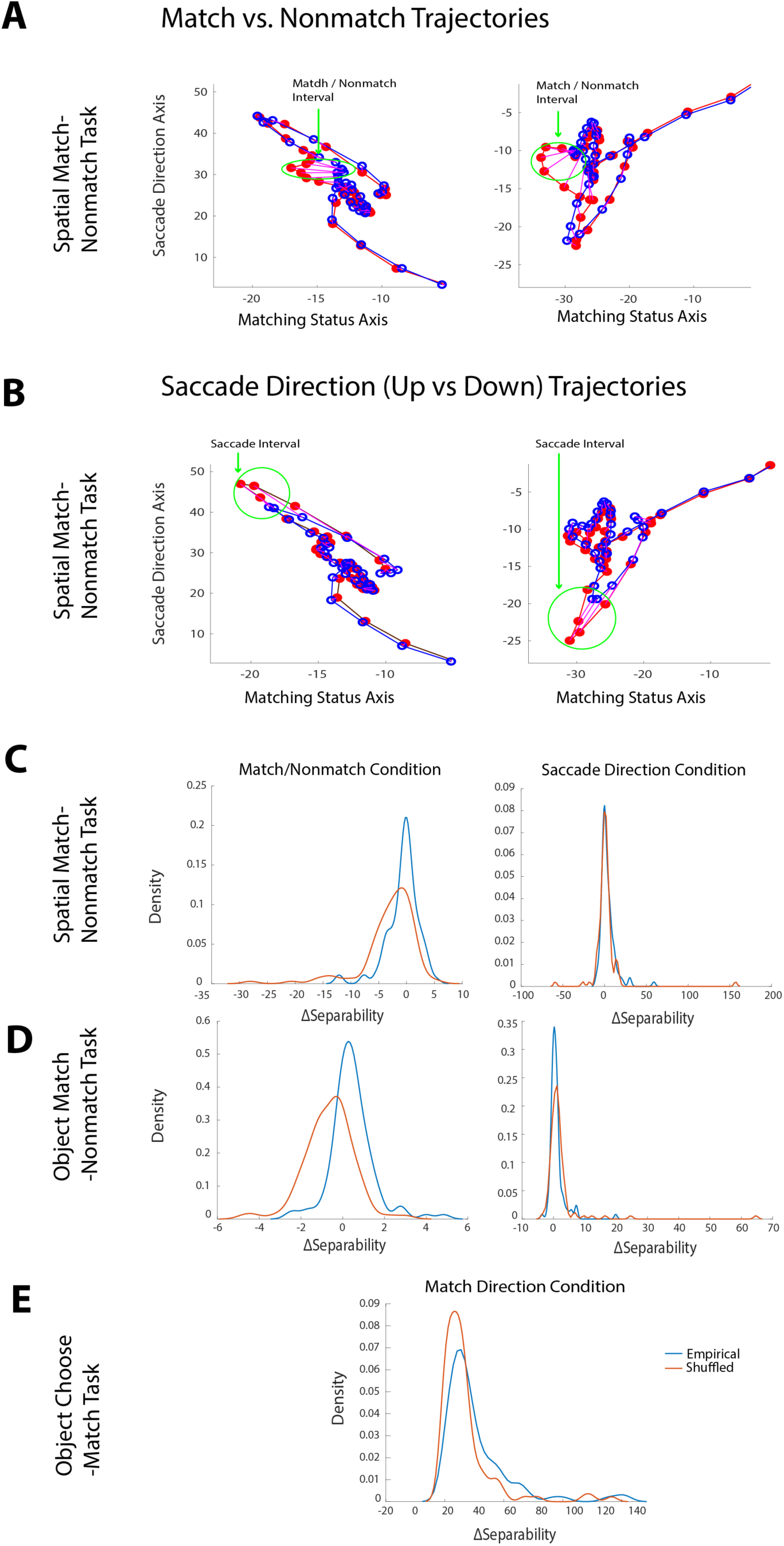
Dynamics of populations responses in PFC. A. Comparison of the complete neural trajectories of novice (left) and expert (right) training across the match (blue) and non-match (red) conditions of the exemplar spatial match-nonmatch task. Separability is compared using the interval that is highlighted in green. This interval was manually determined as the time points that display the greatest distance between conditions. B. Same as Part A for the rewarded target up (blue) and down (red) conditions of the spatial match-nonmatch task. C-E. The distribution (y-axis) of the change in separability (x-axis) between empirical novice and expert neural trajectories (blue) versus a null distribution (orange) that was calculated from a pair of randomly shuffled surrogate datasets. Separability was defined as the Euclidean distance between trajectories across both axes of a two-dimensional subspace capturing the variance due to the combination of the presented stimulus (i.e. match or nonmatch in the spatial and object match-nonmatch tasks, and circle or triangle stimulus presentation in the object choose-match task) and the saccade direction of the rewarded choice target (i.e. up or down) task conditions. This was examined for the match and nonmatch conditions (left) and the target direction conditions (right). Experimental tasks were examined in the order of C. spatial match-nonmatch task, D. object match-nonmatch task, and E. object choose-match task.

We wished to test whether differentiation between task conditions would be represented by increased differences in their respective neural trajectories (Kobak et al., 2016).To perform such tests, we created a distribution of empirical trajectories by subsampling trials from each neuron and creating 100 permutations for the trajectories of the novice and expert training stage. We then calculated a null distribution, wherein all trials from both the novice and expert datasets were pooled together, and then randomly shuffled into two surrogate datasets of equal size to the novice and expert datasets. These surrogate datasets were used to plot firing rate trajectories in a state space defined by the same principal components to compute the null distribution over the course of 100 permutations.

We first sought to compare neural trajectories during the presentation of the match and nonmatch stimuli. In the spatial match-nonmatch task, we first grouped trials into two groups, depending on whether the sample stimulus was a match or nonmatch (see inset of Fig. 4A-left). Each trial group contained trials in which the monkey saccaded either up or down (as illustrated in the inset of Fig. 4B-left). We then examined the separation of neural trajectories in state space specifically during the time of the sample stimulus presentation, in the novice and expert training stages (Fig. 5A and 5B, respectively). For each training stage, we normalized the mean trajectory separation of the trials grouped based on match/nonmatch appearance by dividing this value by the separation of the trials grouped by the saccade-up vs. saccade-down conditions during the sample stimulus time interval. We then compared the trajectory separation in novice and expert data to obtain a value we refer to as ΔSeparability (Fig. 5C-E). A distribution of ΔSeparability values was obtained for the empirical data, by sampling different trials of each neuron (Blue curves in Fig. 5C), and also for the data obtained by randomizing the novice and expert labels (orange curves in Fig. 5C). The empirical increase in separability between the novice and expert training was significantly higher than what would be expected by chance (Wilcoxon test; p<E-4; Fig. 5c-left). This result mirrored the increase in match vs. nonmatch decoding accuracy described above (Fig. 4a - left).

We repeated the same analysis for the separability of neural trajectories in the spatial match/nonmatch task, during the saccade interval (Fig. 5B). We now grouped trials based on saccade direction, examined trajectory separation during the saccade time interval and normalized the separability by the mean distance of trials grouped based on match/nonmatch (Fig. 5C right). In this case, the difference in separability between novice and expert training stages did not reach statistical significance over chance (Wilcoxon test, p=0.088). This result mirrored the difference in decoding accuracy described in Fig. 4B-left.

Importantly, the PCA analysis also revealed an increase in separability in the latent space for the object match-nonmatch task at the time of the sample (match/nonmatch) stimulus presentation (Fig 5d-left, Wilcoxon test, p<E-4). This was so, even though decoding of the match and nonmatch conditions had failed to reveal a clear improvement (in Fig. 4A-center).

In the case of the object choose-match task, we computed match/nonmatch separability at the time the two targets were presented and the monkey needed to choose one of the two. In this case too, separability increased from the novice to the expert training stage (Wilcoxon test, p=0.0005). In conclusion, PCA analysis revealed some common patterns of training effects in latent variables across our tasks, even in cases when explicit decoding of task conditions was not evident.

## DISCUSSION

Prior studies have established that different aspects of working memory such as processing speed, capacity, or the ability to multitask can be modified through training (Klingberg et al., 2002b; Bherer et al., 2008; Constantinidis and Klingberg, 2016). Such training has been proven to be beneficial particularly for clinical populations, such as stroke, ADHD, and schizophrenia patients (Klingberg et al., 2002a; Westerberg et al., 2007; Jaeggi et al., 2008; Subramaniam et al., 2012) as well as aged adults (Anguera et al., 2013). A debate still exists regarding if and how these benefits may generalize (“transfer”) to untrained domains, including everyday life (Cortese et al., 2015; Schwaighofer et al., 2015).

A variety of corresponding changes in neural mechanisms, particularly in the prefrontal cortex—which is widely considered to be the seat of WM—have been described (Qi and Constantinidis, 2012b; Averbeck and O’Doherty, 2022). For example, prior studies have demonstrated how the decoding of task variables from the activity of prefrontal populations has provided a strong neural correlate of rule learning (Meyers et al., 2012). Neurophysiological research—supported by fMRI studies—has also highlighted increases in prefrontal activity after training to indicate increased specialization or strengthened representations of stimuli (Garavan et al., 2000; Mendoza-Halliday and Martinez-Trujillo, 2017). However, other studies have identified decreases in activity after training that may indicate possible improvements in efficiency instead (Schneiders et al., 2011; Takeuchi et al., 2013). Studies have also demonstrated more subtle training effects in firing rate, such as modifications to network modularity and correlation (Qi and Constantinidis, 2012b; Finc et al., 2020). The malleability of WM is thus believed to be mediated by the underlying plasticity in neural responses. Such plasticity has been observed over the course of even individual training sessions, and also during the more extended process that is entailed by rule learning—as shown by our current training paradigm (Asaad et al., 1998; Bartolo and Averbeck, 2020). Rule learning has long been examined through similar reversal tasks as a measure of behavioral flexibility (Schoenbaum et al., 2000; Izquierdo et al., 2017). However, reported results of training are most often limited to an individual task and it has been difficult to determine common and unique elements, investigated with the same experimental and analytical techniques, other than artificial neural network simulations (Masse et al., 2019; Xie et al., 2022).

### Prefrontal activity changes

Prior studies have demonstrated strong evidence of prefrontal persistent activity dynamically incorporating multiple types of both spatial and object information (Amengual and Ben Hamed, 2021; Curtis and Sprague, 2021) with the overall effects of training suggesting increased recruitment of more prefrontal neurons and generation of persistent activity at higher levels (Tang et al., 2022). Our present study was therefore guided by experimental and theoretical predictions to examine whether rule learning could be characterized by changes in firing rate across all tasks. Somewhat unexpectedly, we found that the overall increase in firing rate after training was not a general phenomenon across different tasks. Instead, the progression of rule learning seemed to move firing rate in different directions for different monkeys/tasks, both increasing and decreasing. This result mirrors the conflicting fMRI studies that have suggested the former to indicate strengthened neural representations that underlie short-term information maintenance and manipulation (Garavan et al., 2000; Hempel et al., 2004; Mendoza-Halliday and Martinez-Trujillo, 2017), just as the latter has been suggested to indicate improvements in efficiency (Schneiders et al., 2011; Takeuchi et al., 2013). Our observation of both effects across tasks—increases and decreases—would therefore imply that this debate is not mutually exclusive to one side or the other, and instead, either effect may be responsible for mediating training, as determined by the individual monkey/task.

Nevertheless, at least some common changes in firing rate were evident in our study. For example, our observed trend of a significant decrease in choice epoch activity, particularly early in training, seemed to directly parallel the period of behavioral adjustment wherein each monkey gradually learns the rule—through trial and error—that the rewarded choice target is not random, but instead, is determined by the previously presented stimuli. This was the rule learning that allowed our monkeys to better anticipate which choice would be rewarded in their assigned tasks. Decreased choice epoch activity thus paralleled the observed trend of decreased DIP magnitude that behaviorally characterized the acquisition of the rule until eventually, the rule was mastered and choice epoch firing rate returned to its original state to reflect this new mastery.

The more consistent effect we observed was that of an increase in unexplained variance after training, independent of mean firing rate changes. Prior studies have shown that in real life, animals learn reversal learning tasks via reinforcement learning based on reward outcomes (Kim et al., 2025). In such tasks, increased unexplained variance in firing rate has been observed during reversals due to the encoding of decisions that represent a belief about the current state of the task—which is then contradicted by such reversals (Bartolo and Averbeck, 2020). The change in choice preference reversal that followed from this contradiction is thus predicted by increased unexplained variance as the monkey encoded the task factors that this new preference demanded. Such increases were suggested to be caused by prefrontal neurons dynamically encoding decisions associated with Bayesian subjective values to represent a belief about the current state of the task. To examine how this concept may apply over the process of rule learning, we ran a linear regression to compare how effectively the pre-training model of firing rate could explain variance during novice and expert training based on the three factors of matching status, rewarded target direction, and reward status, this is, whether the trial was answered correctly or not. This revealed a significant increase in unexplained variance across all monkeys/tasks, suggesting that as the monkeys progressively acquired the new rule, they increasingly encoded new task factors that were not linearly related to the three variables of our model. Our effect was attributed solely to less variance being explained during the delivery of reward. In other words, delivery of reward is very informative, and neural activity tracks this variable early in the training when monkeys rely on this feedback to adjust their behavior in each trial but becomes less important when monkeys understand the rule and the outcome becomes more expectable.

### Encoding of new factors

One of the most important issues related to the role of prefrontal activity is how the representation of different types of information may be altered by rule learning to achieve more effective task performance (Rigotti et al., 2013). We applied a decoding analysis to examine how training affects the decoding of different types of task factors, with the prediction that training would lead to an overall improvement in decoder performance across tasks—and by extension, an improved representation of the information needed for success. Somewhat unexpectedly, each of our tasks revealed a different pattern of decoding, with only the spatial match-nonmatch task displaying the improved decoding of match/nonmatch information that might have been expected. When we tested the two object tasks to determine if the same pattern would occur, the object match-nonmatch task revealed no clear training effects, while the object choose-match task revealed a strong bias in neural activity, early, which disappeared as training progressed.

Yet when we used PCA analysis to examine latent population dynamics (Mante et al., 2013; Mozumder and Constantinidis, 2023), we found an increased separation of population response trajectories in state space during the match/nonmatch presentation across all tasks. This suggests that rule learning leads to at least a subtle differentiation between task conditions in prefrontal activity that was not evident from the decoder analysis which examined each explicit task variable separately.

### Confronting the object-spatial divide

Our findings demonstrate that although both spatial and object information may be represented in prefrontal activity, these modalities seem to present a variety of key differences that would imply at least some degree of task-specificity in the neural mechanisms of rule learning (Nystrom et al., 2000; Ester et al., 2009). Most importantly, a large proportion of neurons represented spatial location, were modulated by the task, and differentiated between a left and right stimulus, when these represented a match or nonmatch condition. In contrast, few neurons represented the object stimuli, both before and after training, resulting in a relative paucity of information about object shape when it signaled a match or nonmatch.

Such differences are emphasized by studies that identified dynamic interactions with lower cortices, such as the primary visual cortex, during training of object WM tasks as well (Fyall et al., 2017; Skirzewski et al., 2021). Any debate on the generalization of mechanisms across spatial and object rule learning should not be seen as an all or nothing divide. Instead, both theories seem to work in tandem to an extent that is determined by context.

There is also some evidence of prefrontal persistent activity dynamically incorporating multiple types of both spatial and object information, thus implying that this may not be an exclusive mechanism to spatial WM alone (Amengual and Ben Hamed, 2021). However, the extent to which this may overlap between modalities is highly debated, and a lack of shared training effects in our tasks would imply a separation of spatial and object rule learning mechanisms, consistent with prior theories that object stimuli are maintained via lower cortices—such as the primary visual cortex—while the PFC plays a more supervisory role instead (Ester et al., 2009; Lawrence et al., 2018).

Our current training paradigm would be unable to fully assess the potential influence of phenomena such as match suppression, where activity elicited after repeated presentation of the same stimulus is reduced, which may be informative regarding the matching status of a given stimulus with a prior stimulus (Grill-Spector et al., 2006). Identifying which factors determine when a given rule learning mechanism is either generalizable or task-specific may also provide further explanation for previously nebulous phenomena, such as the previously noted “special status” of spatial information in WM via the bump attractor model (Foster et al., 2017). Furthermore, such investigations will allow new insights on the overall role of persistent activity in the PFC and how the generalization of this mechanism may compare to the generalization of alternative mechanisms that have been proposed for WM, such as the modulation of synaptic resources (Mongillo et al., 2008). To that end, our results provide a framework for exploring how prefrontal mechanisms parallel the progression of rule learning, ultimately laying the foundation for addressing each of these questions and beyond.

## METHODS

### Test Subjects

Data obtained from four rhesus monkeys (*Macaca mulatta*, ages 5-9 years, weighing 5-12kg, three male and one female), were analyzed in this study. None of these animals had any prior experimentation experience at the onset of our study. Monkeys were single housed in communal rooms with sensory interactions with other monkeys. Access to water was restricted during training to motivate task performance. All experimental procedures followed guidelines set by the US Public Health Service Policy on Humane Care and Use of Laboratory Animals and the National Research Council’s Guide for the Care and Use of Laboratory Animals as reviewed and approved by the IACUC (Institutional animal care and use committee) of Wake Forest University and/or Vanderbilt University.

### Experimental Setup

In all three of our experimental tasks, monkeys sat with their heads fixed in a primate chair while viewing a monitor positioned 68 cm away from their eyes with dim ambient illumination and were required to fixate on a white square of 0.2 degrees appearing in the center of the screen. During each trial of every task, the monkeys were required to maintain fixation on the square while visual stimuli were presented either at a peripheral location or over the fovea, to receive a juice reward. Any break of fixation immediately terminated the trial, and no reward was given.

Eye position as monitored in the spatial match-nonmatch task via a non-invasive, infra-red eye position scanning system (model RK-716; ISCAN, Burlington, MA). Eye position was sampled at 240 Hz, digitized, and recorded. Similarly, eye position was also monitored in both object tasks via another non-invasive, infra-red eye position scanning system (model ETL-200; ISCAN), sampled at 500 Hz, digitized, and recorded. The system achieved a <0.3° resolution around the center of vision. The visual stimulus display, monitoring of eye position, and synchronization of stimuli with neurophysiological data were performed with in-house software implemented on the MATLAB environment (The MathWorks), using the Psychophysics Toolbox (Meyer and Constantinidis, 2005).

### Match-Nonmatch Tasks and Pre-Training Paradigm

Three monkeys (Subjects NI, MA, and FI) were trained in either a spatial or object version of a match-nonmatch task based on the multi-phase training paradigm of a prior study (Tang et al., 2022). This previous multi-phase training paradigm consisted of four phases that were designed to ensure that specific task elements would be acquired in sequence (Fig. S1). First, the monkey was presented with two stimuli in rapid succession and had to indicate if they appeared at the same or different locations by selecting one of two choice targets. During this phase, daily training sessions involved the presentation of a cue stimulus at the right of the fixation point followed by a sample stimulus appearing at either a matching location (right) or a non-matching location (left), on different days (Fig. S1b). At this stage, the monkey could simply examine the choice targets, determine which one was rewarded during the block, and repeatedly select it in following trials. In the second phase, the monkey was presented with alternating blocks of match and nonmatch trials, of decreasing block length, until they were randomly interleaved, requiring the monkey to gradually learn to associate the second stimulus location with the corresponding choice target (Fig. S1c). Examining the rule learning that occurred over the course of this second phase formed the foundation of our current experiment (Fig. 1a). However, the previous multi-phase training paradigm also included a third phase wherein the monkeys had to generalize the task to new stimulus locations, appearing at a 3 × 3 grid (Fig. S1d), as well as a fourth phase, wherein the delay period was increased to place a greater demand on WM (Fig. S1e).

Our current training paradigm thus utilized the same spatial match-nonmatch task from the second phase of this previous training paradigm, as well as a parallel object version of this match-nonmatch task that was created specifically for the needs of our current experiment. In either case, our monkeys were given a window of 2 seconds to begin fixation, or the trial would be automatically aborted. Following this initial acquisition of the fixation point, the trial would begin with a fixation epoch in which no stimuli would be shown (1.0 s). This was followed by the presentation of a single stimulus that was always the same and was referred to as the cue (0.5 s). This first stimulus was always a square that appeared to the right of fixation in the spatial task, and always a circle that appeared over the fixation in the object task. This was followed by a very brief delay epoch (0.25 s) and then a second stimulus presentation, known as the sample, which could vary in either location (appearing to the left or right of fixation in the spatial task), or shape (appearing as a triangle or circle in the object task). After the second delay epoch (0.25 s), the two choice targets (Diamond and H-shape for the spatial task, Green and Blue for the object task) appeared with the fixation point turning off, either above or below the fixation point, but randomly switching between trials (0.5 s). Only one of these two choice targets would produce reward in each trial, and this was determined by the cue and sample stimuli that were previously presented. If these stimuli matched each other, the “Diamond” and “Green” choice targets were rewarded. Likewise, if these stimuli did not match, the “H-shape” and “Blue” targets were rewarded instead (Fig. 1a).

At the start of our experiment, all monkeys were already able to maintain fixation and had been exposed to the visual stimuli that would eventually be incorporated into their respective tasks. Moreover, monkeys had been taught a pre-training version of the spatial and object match-nonmatch task, wherein sessions would present trials that used only one of the two possible stimulus combinations, without any trials that used the other stimulus combination. The presented stimulus combination that was selected for any single training session would alternate between the two options over the course of multiple sessions but would not mix. This meant that the single rewarded choice target remained constant throughout the span of the entire training session and thus taught the monkeys to make an eye movement to one of two choice targets to determine that only one of them would be rewarded. Then, once the monkey learned which choice target would be rewarded for the session, they could simply repeatedly select that target for the rest of the trials in that session to continue receiving reward. This discretized pre-training paradigm ensured that different task elements would be acquired in sequence. Specifically, monkeys had no need to pay attention to which of the two possible cue and sample stimulus combinations were being used in each trial and were thus not yet aware of the difference between trials with matching or nonmatching stimuli.

Our current training paradigm was designed to introduce a new task element by progressively interweaving the two possible stimulus combinations into the same training sessions. This was carried out by requiring for the monkeys to periodically switch between trials with matching and nonmatching stimuli after a set block of correct trials, the size of which was constant and manually predetermined at the start of each session. As a result, the rewarded choice target would change with each trial block, and the monkeys had to learn to identify the difference between trials with matching or nonmatching stimuli, since the presented stimulus combination would be responsible for determining which choice target would be rewarded at the end of each trial. During the first training sessions when the monkey had not yet begun to learn the new rules of this experiment, they were expected to perseverate in their behavior, and as a result, they would almost always fail the trial in which there was a reversal between matching and nonmatching stimuli. This led to a consistent Drop In Performance (henceforth abbreviated as DIP) at the reversal trial, and as a result, the shorter these trial blocks became before a reversal between the two possible stimulus combinations, the more the monkeys would be forced to adapt their behavior to avoid failure.

### Object Choose-Match Task

Two monkeys (Subjects IT and FI) were trained in an additional object working memory task, the object choose-match task as well. As with the spatial and object match-nonmatch tasks, each trial began with a fixation epoch in which no stimuli were shown (1.0 s). This was followed by a cue stimulus (either a circle or a triangle) that was displayed at the center of the screen (0.5 s), and then a delay epoch (0.25 s), after which two stimuli (both a circle and a triangle) were displayed on diametrically opposite sides of the screen (0.5 s), prompting the monkeys to indicate which of these matched the first presented stimulus via an eye movement. These trials blocks alternated which object was used as the initial cue stimulus, thus allowing the learning of the same rule from the spatial and object match-nonmatch tasks—that when trials changed between presenting different stimuli, the rewarded choice target would be change as well—to be examined in a different context as well (Fig 1a-right).

### Behavioral Analysis

For all three tasks, performance was calculated for each session by dividing the number of successful trials by the total completed trials, excluding any trials in which fixation was broken. Each monkey was assessed individually to ensure that a sufficient level of task performance (>60%) was maintained throughout the course of this experiment (Fig. 1b and Fig. S2a). Our training paradigm did not allow subsequent sessions to progress with the decrease in trial block size unless the monkey proved able to consistently maintain sufficient performance.

### Behavioral Strategy Analysis

To quantify the behavioral strategies employed by each monkey across training sessions, we modeled each trial as arising from one of five candidate strategies and estimated the relative contribution of each strategy on a per-session basis. Sessions with fewer than 100 completed trials were excluded from analysis.

Five candidate strategies were defined, each making a deterministic or probabilistic prediction about the probability of choosing the upward saccade target on each trial: 1. Random choice: The monkey saccades upward or downward with equal probability (p = 0.5) on every trial, regardless of trial history or stimulus information. 2. Preferred location: The monkey consistently saccades to the same spatial location on every trial. 3. Preferred color: The monkey consistently chooses based on the color of the choice target, regardless of the matching status of the stimuli. 4. Win-stay-lose-shift: The monkey repeats the color choice that was rewarded on the previous trial (win-stay) and switches to the alternative color after an unrewarded trial (lose-shift). On the first trial of a session, when no prior outcome was available, this strategy assigned p(up) = 0.5. 5. Match-nonmatch rule: The monkey uses the correct task rule, choosing the target color associated with the match or nonmatch status of the current trial.

On each trial i, each strategy s generated a prediction for the probability of choosing the upward saccade, denoted pₛ(up | trial i). If the monkey chose upward on that trial, the likelihood assigned to that strategy was pₛ(up); if the monkey chose downward, it was 1 − pₛ(up). These raw likelihoods were normalized across strategies on each trial to sum to 1, yielding a per-trial weight for each strategy:

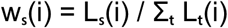

where Lₛ(i) is the likelihood of the observed choice under strategy s on trial i, and the sum in the denominator runs over all five strategies. The session-level weight index for each strategy was then computed as the mean of these per-trial weights across the first 80% of trials in the session:

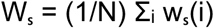

where N is the number of trials used. This normalization ensures that the five weight indices sum to 1 for each session, allowing them to be interpreted as the relative probability that each strategy best accounts for the monkey’s behavior during that session. Sessions in which the animal was excluded due to insufficient trial counts were assigned weight indices of zero and excluded from all subsequent statistical comparisons.

### Implant Surgery

All subjects were initially acclimated with the laboratory and trained to maintain fixation on a white dot while visual stimuli appeared on the screen. After this initial stage of training was complete, monkeys were implanted with an MRI-compatible headpost device. Another surgery was used to implant a chronic array of electrodes in their lateral PFCs (Fig. S3a). All cases of surgery were carried out under inhalant anesthesia. Opioid analgesics were administered after the surgery, and the animals were allowed to recover for at least 3 weeks before behavioral sessions began anew.

For the two monkeys assigned to the spatial task (Subjects NI and MA), the chronic implant was designed in-house and encompassed 64 parylene-c insulated, Iridium electrodes that tapered from a 40 mm diameter to an exposed electrode tip that ranged from 5–7 mm long (Spingath et al., 2011). The implant comprised an 8 × 8 grid of electrodes, with adjacent electrodes spaced 0.75 mm apart from each other, thus covering an area of 5.25 mm × 5.25 mm. This electrode array targeted the dorsolateral PFC, with electrode tracks descending in both banks of the principal sulcus.

For the two monkeys assigned to the object tasks (Subjects FI and IT), the chronic implant was a chronic 128 Grey Matter Research electrode array. This implant comprised an (incomplete) 12x12 grid of tungsten electrodes with a 250 μm diameter, spaced 1.5 mm apart from each other. The electrode array targeted the dorsolateral and ventrolateral PFC, centered across the dorso-ventral and anterior-posterior axis. For the purposes of our analysis, the anterior-posterior axis was defined as the line connecting the genu of the arcuate sulcus to the frontal pole. The recording coordinates of each neuron was projected onto this line, with position expressed as a proportion of the line’s length.

Across all monkeys, each electrode used for recording had a 1 MΩ impedance at 1 kHz. The position of each electrode array was determined based on MRI and then verified during the stereotaxic implantation surgery. Electrodes from both implants could be advanced into the cortex independently of each other and electrode depths were adjusted as needed to optimize placements, up to 5 mm (in the banks of sulci).

### Recording Neural Activity

Once electrode positioning was finalized and the monkeys recovered from surgery, task training and neuro-physiological recordings from the array commenced. Neuronal data from each electrode were recorded over the course of training using an unbiased spike selection procedure, with no regard to the response properties of the isolated neurons. For all monkeys, the signal from each electrode was amplified and sampled continuously at 30kHz and stored for off-line analysis. Data was band-pass filtered between 500 Hz and 8 kHz for spike identification.

The two monkeys assigned to the spatial task (Subjects N and M), used a Cerberus system (Blackrock Microsystems, Salt Lake City, UT), with the threshold for spike acquisition was set at 3.5 × RMS of the baseline signal, for each electrode, each day. The two monkeys assigned to the object tasks (Subjects F and I), by contrast, used a modular data acquisition system (OpenEphys system, FHC, Bowdoin, ME). Semi-automated cluster analysis was carried out through Kilosort, applying principal component analysis of the waveforms to sort recorded spike waveforms into separate units.

To ensure a stable firing rate in the analyzed recordings, any neurons with a mean Fano factor > 2 were identified. As with our prior studies, we examined Fano factor via the methods of Churchland and colleagues, using the algorithm developed by these authors and made available online at: http://www.stanford.edu/shenoy/GroupCodePacks.htm (Churchland et al., 2010; Qi and Constantinidis, 2012b). Data for each neuron was initially treated separately. Spike counts were computed in a 100-ms sliding window moving in 20-ms steps. The method computed the variance and mean of the spike count across trials in each time bin and performed a regression of the variance to the mean. This slope of this regression represents the Fano Factor reported at the given time bin, and any trials from high-variability (mean Fano factor > 2) neurons that displayed an outlier level of activity relative to the total activity of that neuron were filtered out (0.7% of cells from the spatial task, 16% of cells from the object match-nonmatch task, and 3.6% of cells from the object choose-match task). More specifically, these outlier levels of activity from high-variability neurons were identified by fitting a two-component mixture model (Poisson + Gaussian) to each of the given neuron’s trial activity, using a Bayesian information criterion (BIC) to determine if the distribution is genuinely multimodal, calculating the separation between modes (i.e. peaks in distribution), and then finally, removing all trials from the smaller modes. Each high-variability neuron was examined individually to identify and remove these outlier trials.

### Firing Rate Analysis

Analyses of neural firing rate were implemented with the MATLAB computational environment (Mathworks 2019, Natick, MA). Firing rates were assessed separately for each epoch across all cells. This was also repeated for each epoch after subtracting the mean baseline fixation firing rate, thus isolating any activity changes that were caused by the task itself, rather than spontaneous noise (He, 2013). Only correct trials were applied in these analyses to reduce possible confounds and to focus on when the monkey’s behavior would be most reflective of the rule learning that would be needed for optimized performance.

To begin addressing our question of how parallel activity-based mechanisms may be applied across spatial and object rule learning, we applied a series of pairwise Pearson Correlations, based on what has been used in prior studies (Salinas and Sejnowski, 2000). In this case, we were examining the mean correlation coefficient that was taken from all combinations of monkey pairs to determine the degree to which each firing rate variable may predict rule learning for all monkeys/tasks over the course of the novice/expert training stages, with the latter split into mid-expert and late-expert for this analysis. The transition from mid-expert to late-expert was defined as the sessions that occurred after asymptotic performance had been achieved. No quantitative measure was perfectly monotonic across all monkeys, so a qualitative measure was used instead.

We thus created a matrix C, which had X columns by Y rows, where X represented each monkey, and Y represented the novice/expert training stages, with the latter split into mid-expert and late-expert. Each entry consisted of the mean of a given firing rate variable for the given monkey and training stage. The correlation coefficient of our sampled neurons, r, was thus calculated through the following equation:

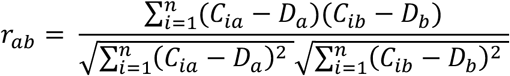

Here, r represents the correlation coefficient between a single pair of monkeys, a and b. Likewise, D represents the means of the firing rate variable across training that was drawn from this pair of monkeys. By examining the mean correlation coefficient that was drawn from all pairs of monkeys, repeated separately for each firing rate variable, we were able to determine which of these firing rate variables had the strongest linear relationship with training across monkeys.

### Generalized Additive Mixed Models

Given that our hypothesized relationship between training and prefrontal activity may not necessarily be captured through linear methods alone, we applied a series of generalized additive mixed models (GAMMs) as a more sensitive method of quantification. GAMMs are a flexible, semiparametric method for identifying and estimating nonlinear effects of covariates on the outcome variable when observations are not independent. We therefore predicted that this would reveal possible nonlinear relationships between training and prefrontal activity. To this end, we examined how the single predictor variable of DIP magnitude may affect the outcome variable of firing rate over the course of training based upon a prior study from our a Zhu et al., 2024):

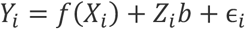

In this equation, Y_i_ represents epoch firing rate for the ith observation, and X_i_ represents the DIP value for that observation. The relationships between each predictor variable and the response variable are represented by *f*, which is a spline with penalty, meaning that smoothing of the curve has been enforced using up to a maximum of five basis functions for the sake of preventing overfitting. Moreover, Z_i_b represents the random effects in the model, where Z_i_ is a matrix that specifies the random effects design for the ith observation, and b is a vector of random effects coefficients. Finally, ɛ_i_ represents the error term for the ith observation.

Just like our previous firing rate analyses, the firing rates variables that were used in our GAMM analyses were assessed separately for each epoch across all cells, and also across all cells with the mean baseline fixation epoch firing rate was subtracted from the other epochs.

Moreover, we once again relied exclusively on correct trials to reduce possible confounds and to focus on when the monkey’s behavior would be most reflective of the rule learning that would be needed for optimized performance.

GAMM analyses were implemented using the mgcv package for R with a smooth function of DIP across sessions as a covariate, using a thin plate regression spline basis to estimate this smooth function. This allowed us to model nonlinear effects using a flexible, automatically adaptive curve with theoretically optimal smoothness control, with each datapoint used to construct our splines automatically in lieu of manual knot selection.

Random effects included the subject-specific intercepts and slopes for DIP. The outcome of firing rate—which was continuous—was examined via a Gaussian family and identity link function. For those models for which there was a statistically significant fixed effect, the gratia package was used to conduct exploratory post-hoc analyses to identify significant training effects by approximating the derivatives of each estimated smooth function of DIP via the method of finite differences, and a simultaneous 95% confidence, which were calculated by randomizing the labels of different monkeys at each x-axis point (representing mean session DIP) and using that to refit the GAMMs over 100 repetitions.

### Unexplained Variance in Firing Rate

To confront the question of how training may affect the computation efficiency of encoding task relevant stimuli—especially during the reversal between trials with different stimuli—we analyzed changes in firing rate variance that could not be explained by the task factors of the presented stimulus, the direction of the rewarded choice target, and whether the trial was answered correctly or not, examined across trials based on their position relative to the reversal itself. This analysis thus relied upon both correct and error trials, as both were necessary to assess the given task factors, and was evaluated through a linear regression model with the following equation:

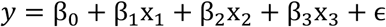

Here, y represents the dependent variable of firing rate, modeled as a function of β0, which represents the baseline firing rate as an intercept term, and x1, x2, and x3, which represent the predictors (independent variables of the three examined task factors). These three factors were the presented stimulus, the direction of the rewarded choice target, and whether the trial was answered correctly or not, examined across trials based on their position relative to the reversal itself. Next, β1, β2, and β3 represent the regression coefficients of these task factors and quantify how much each predictor contributes to explaining the firing rate as slopes that show how much the firing rate changes per unit changes with each predictor. Finally, ɛ represents the error term (residuals), which define the unexplained variance as the difference between the observed firing rate and the model’s prediction. To fit the model, spike count data were generated using a moving window of 400ms wide and 100ms steps, from 1.5s before cue onset to 2.5s after in the case of the spatial and object match-nonmatch tasks, or 1.5s before cue onset to 1.5s after in the case of the object choose-match task. The linear model was then fitted to each time bin individually, using all possible trials from each distance relative to the reversal trial. The percentage of unexplained variance ratio (SSE/SST) was then calculated for each relative distance to the reversal, and max-normalized across all distances (i.e. the trial before, during, and after the reversal). Lastly, the results across all time points were averaged for each relative distance to the reversal in each cell. This unexplained variance was then examined across relative distances to the reversal and across novice/expert training through a series of two-way ANOVAs. Only cells with at least three trials at each of the examined distances relative to the reversal were included in this analysis.

### Decoding of Task Variables

To understand how training affects the types of information represented in neural populations, a new set of decoder analyses were added using the supervised max-margin model of a support vector machine (SVM) on the MATLAB environment with a linear kernel function, based upon what has been used in our prior studies of the PFC (Dang et al., 2021). We predicted—just as training was observed to shift firing rate in different directions for different tasks/monkeys—training may shift the separation of task factor conditions in the resulting max-margin boundary (i.e. the hyperplane) in different directions for different tasks/monkeys as well. This would be expressed through different changes in decoder performance for different tasks/monkeys, ultimately providing new insights on the implications of such diverse effects that could not be inferred from firing rate alone. This led us to apply the fitcecoc function to produce a multiclass, error-correcting output codes (ECOC) model using the predictors that were drawn from our recorded neuronal data.

To obtain the predictor data of a single trial, we created a 3-D matrix that had the dimensions of cell number, by the status of the examined task factor, by the total number of 400 ms time bins. The total number of time bins was decided by using a 400 ms sliding window, with a 100 ms step size (with overlap) spanning the full length of the trial (3s for both of the spatial and object match-nonmatch tasks, or 2.25s for the object choose-match task). This resulted in a total of 27 time bins for the spatial and object match-nonmatch tasks and 19 time bins for the object choose-match task. Each entry of the resulting 3-D matrix consisted of the spike count of the individual neuron in the selected time bin and trial’s condition of the examined task variable. This was then used to create a pseudo-population that would provide training data, thus allowing us to test the resulting classifier on the factors of the presented stimulus, the direction of the rewarded choice target (i.e. whether the rewarded target appeared up or down), the configuration of the choice targets (i.e. whether a specific target appeared up or down regardless of whether it was rewarded), the combination of the presented stimulus with the direction of the rewarded choice target, the monkey’s choice of saccade direction, and the monkey’s choice of saccade to a match, via the class label input, that is, a scalar variable indicating the task variable that the decoder was designed to examine. This analysis relied upon both correct and error trials, examining the same number of cells at each stage of training.

Ten-fold cross validation then split our training data into 10 sets, 9 of which were used for training, while only the last tenth of the data was applied for testing, thus producing a cross-validated model that could be used to make a prediction of the data at each time point. Training and testing were first carried out using correct trials only, and then again, separately, using correct trials combined with error trials, to result in two cross-validated models that would be examined independently. We reasoned that the encoding of information does not necessarily imply the utilization of such information, and so the inclusion of error trials in this latter cross-validated model would allow us to dissociate behavioral and sensory encoding, thereby revealing possible changes in the encoding of different types of task data that would not be evident from the use of correct trials alone.

This process was then repeated 10 times, thus allowing all data to be used as both training and test data. The average accuracy of these 10 repetitions was counted as the decoding accuracy for a single iteration, and this entire process was then repeated across 10 iterations, with 95% confidence intervals calculated via linear interpolation on these 10 iterations.

Finally, to determine whether the change in decoder performance over the course training was consistently within chance level, the average decoding accuracy across all iterations was compared across novice vs. expert training at each timepoint through a nonparametric randomization test—also known as a permutation test—wherein all trials from both the novice and expert datasets were pooled together, and then randomly shuffled into two surrogate datasets of equal size to the novice and expert datasets. These surrogate datasets were used to produce two cross-validated models in the same manner as the original decoders, with the difference in their performance at each timepoint being used to compute a null distribution over the course of 50 permutations. We then compared the difference in performance from the original decoders to this null distribution and identified significance at any timepoints where this observed performance surpassed the top 1% of the null distribution (p < 0.04, two-tailed).

### Targeted Dimensionality Reduction (TDR)

To characterize how task-relevant variables—specifically matching status (cue shape in the choose-match task) and the saccade direction of the rewarded target—are represented in the neural population, we employed targeted dimensionality reduction (TDR). This method identifies low-dimensional axes within the neural state space that are maximally sensitive to specific task variables while remaining orthogonal to one another. The neural response matrix R (units × time bins × trials) was first organized by experimental conditions. We defined conditions based on the unique combinations of the two examined task variables: matching status (cue shape in the choose-match task) and choice saccade direction. For each unit, the trial-averaged response X_c_(t) was calculated for each condition c. These condition-averaged responses were then z-scored across all time points and conditions to ensure units with high firing rates did not disproportionately dominate the population analysis.

To focus our analysis on the most prominent features of the population response and reduce trial-to-trial noise, we performed Principal Component Analysis (PCA) on the condition-averaged data. We constructed a data matrix by concatenating the z-scored responses across all conditions and time points. The first Np principal components (capped at 12 or the numerical rank of the data) were retained to define a “denoising” subspace. We defined a denoising operator D as: D = V * V’, where V is the matrix containing the first Np eigenvectors (loadings). This operator was used to project subsequent regression vectors back into the primary task-related subspace.

To identify the contribution of each task variable to the neural activity, we performed a trial-by-trial linear regression for each unit i and each time bin t: r_i_(t, k) = β_match_(i, t) * var1(k) + β_sac_(i, t) * var2(k) + β_0_(i, t) + ε, where r_i_(t, k) is the z-scored response of unit i at time t for trial k, and var1, var2 represent the matching status (cue shape in the choose-match task) and saccade direction variables, respectively. This yielded time-varying regression vectors, β_match_(t) and β_sac_(t), which point in the direction of the state space that captures the variance associated with each variable.

For both variables regression vectors, we first denoised them by projecting them into the denoise PCA subspace: β_denoised_(t) = D * β(t). Then for each variable v, we identified a single time-independent axis by selecting the vector β_denoised_(t) at the time point t where its magnitude (norm) was maximal. To ensure that the resulting task axes were independent, we applied a QR decomposition to the set of regression vectors: [ω_match, ω_sac] = QR([β_match(t), β_sac(t)]). This resulted in an orthonormal basis where each axis represents a specific task variable, purified of contributions from the other. Finally, the condition-averaged population activity was projected onto these orthonormal axes.

Neural data was visualized in the space defined by two task variable related principal components defined as described above, using a 400 ms wide moving window average with a 100 ms step size. Separability was defined as the Euclidean distance between trajectories across both axes of a two-dimensional subspace capturing the variance due to the combination of matching status (cue shape in the choose-match task) and the saccade direction of the rewarded choice target (i.e. up or down) task conditions. To quantify the separability of neural trajectories along each task-relevant axis, we computed a normalized Euclidean distance metric for each dimension separately (matching status and saccade direction). We identified four trial conditions defined by the crossing of matching status (match/non-match) and saccade direction (up/down): match-up (c₁), match-down (c₂), non-match-up (c₃), and non-match-down (c₄). For each resampling iteration, the neural state for each condition was estimated by averaging the projected coordinates across time points 23–24, corresponding to the late sample period. Match and non-match centroids were then computed by averaging across saccade directions within each matching category:

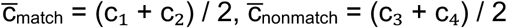

The separability index was then defined as:

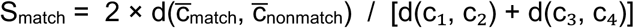

where d(·,·) denotes Euclidean distance in the 2D PCA subspace. This normalization by the mean within-group spread ensures the index reflects between-group separation relative to within-group variability, yielding a value greater than 1 when between-group distance exceeds within-group spread. Separability across the matching status conditions was examined through the interval of the second stimulus presentation (cue presentation in the choose-match task), while separability across the target direction conditions was examined through the interval of the choice target presentation.

The difference in separability between conditions was then assessed across novice vs expert training by calculating a null distribution using the same methods as the decoder analysis, wherein all trials from both the novice and expert datasets were pooled together, and then randomly shuffled into two surrogate datasets of equal size to the novice and expert datasets. These surrogate datasets were used to plot firing rate trajectories in state space corresponding defined by the same two principal components to compute a null distribution over the course of 100 permutations.

We then applied a Wilcoxon rank sum test to compare the vector of all empirical differences between the observed novice and expert neural trajectories against a null distribution that was drawn from the shuffled differences. Empirical and null distributions were drawn by resampling 100 times for each cell and task condition. We then calculated a test statistic based on how many times the empirical values exceed the permutation values. Under the null hypothesis (no difference between groups), empirical differences would be randomly distributed, and so this allowed us to determine whether the observed empirical difference between novice and expert trajectory distances could not plausibly arise from random sampling.

## ACKNOWLEDGEMENTS

Supported by NIH award number R01 EY017077, and F31 EY035546. We wish to thank Chrissy Suell, Kayla Yetman, Kris Clifft, and Jaela Bills for technical help and Emilio Salinas, Terry Stanford, Ben Rowland and Robert Hampson for helpful comments on an earlier version of this manuscript.

## AUTHOR CONTRIBUTIONS

CC designed the experiment. RJ, WD, TG, and JZ performed data analyses. CC, WD, and RJ wrote the paper.

## DATA AVAILABILITY

Data for the current study will be made available through Zenodo upon acceptance of the paper.

## CODE AVAILABILITY

The code used to process the results and generate the figures will be made available at Github upon acceptance of the paper.

## SUPPLEMENTARY INFORMATION

**Supplementary Figure S1:**
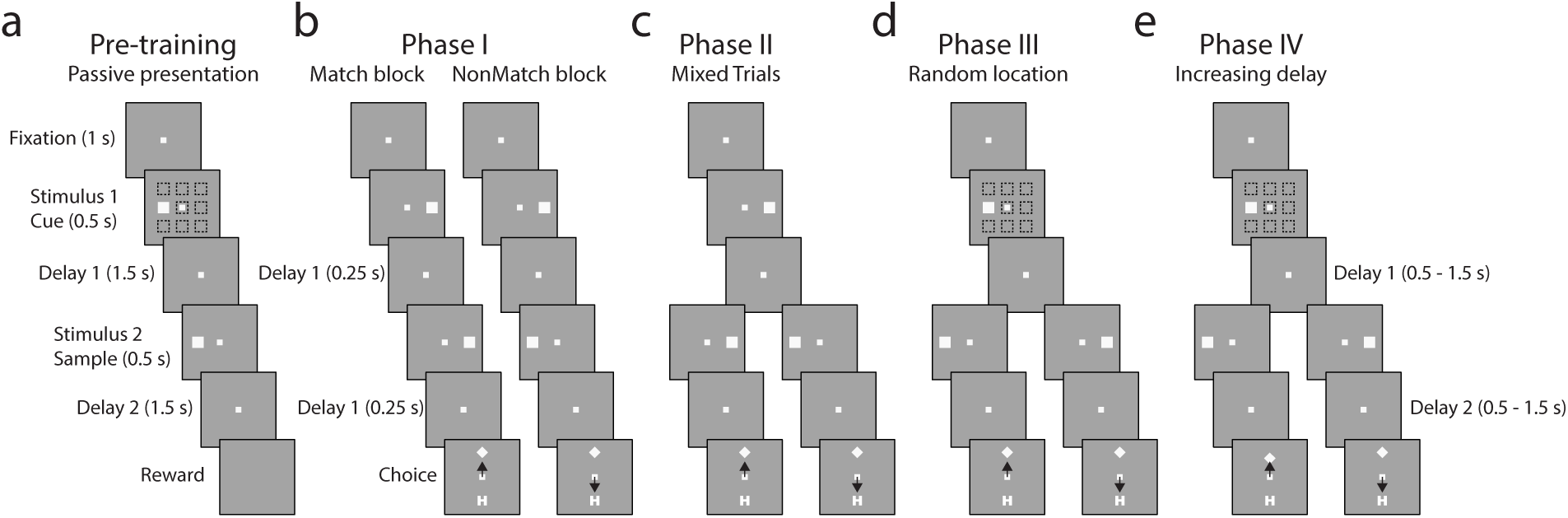
Full training paradigm. A schematic of all training phases from the previous multi-phase spatial match-nonmatch task training paradigm that our current training paradigm was founded on (Tang et al., 2022). A-E Successive frames illustrate the sequence of events in the tasks used in progressing training phases. A. During the passive pre-training phase, the monkey only had to fixate while the stimuli were displayed at any one of the nine locations on the screen. B. In Phase I, a stimulus was always presented to the right, followed by a match stimulus in a block of trials and by a nonmatch stimulus in another block of trials. At the end of the trial, two choice targets appeared, and the monkey had to choose the “Diamond” target in match blocks and the “H” target in nonmatch blocks to get a reward. C. In Phase II, match and nonmatch trials were mixed in a block. This was identical to the spatial match-nonmatch task that was used in our current experiment. D. In Phase III, the stimulus location of the first stimulus could vary. E. In Phase IV, the duration of the delay period increased. The passive stimulus set continued to be presented at the beginning of each session throughout all four phases of training in this paradigm.

**Supplementary Figure S2:**
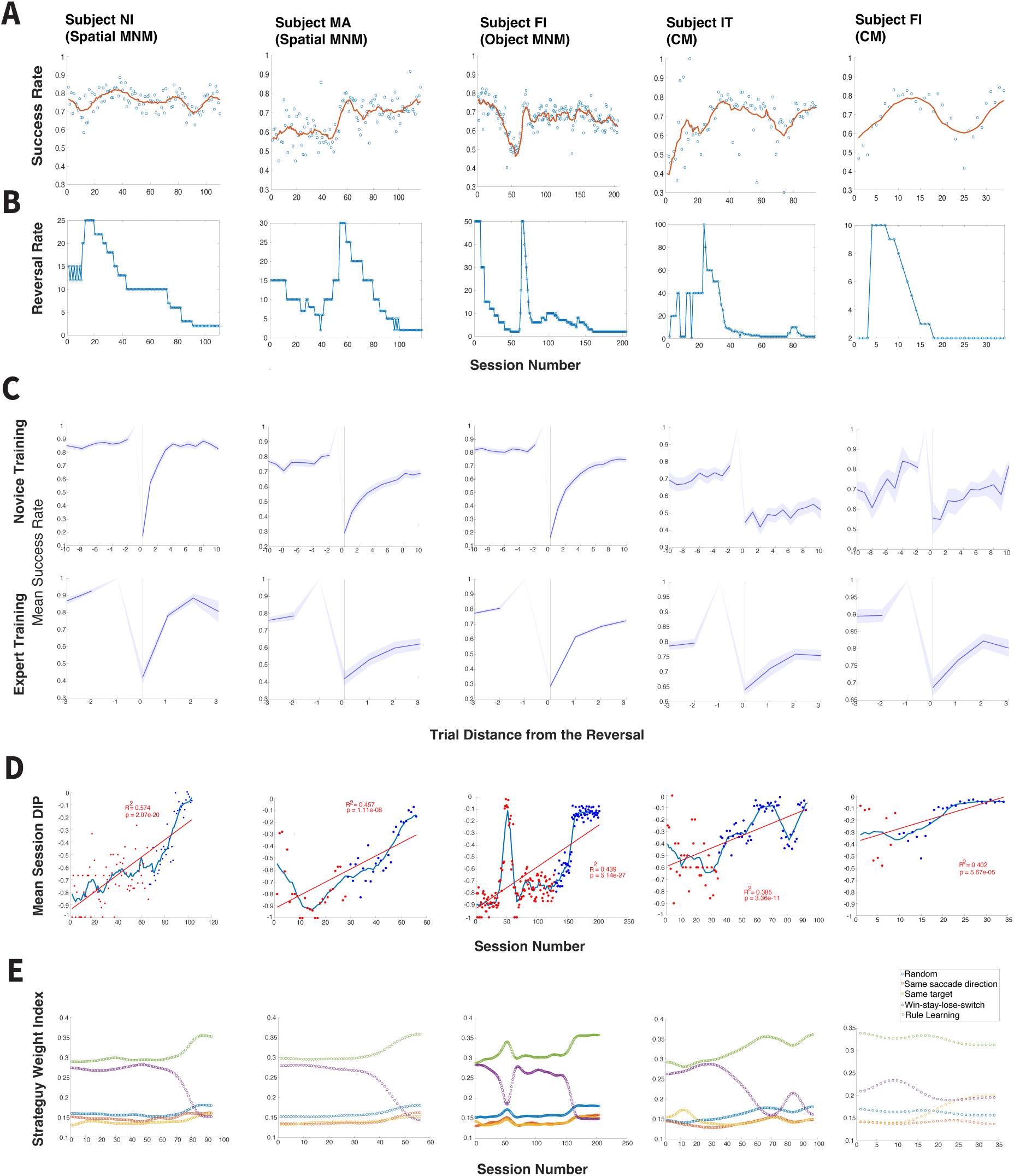
Expanded task behavior. Expanded behavioral results. This includes the mean success rate (A), the reversal rate (B) between the presentation of match and nonmatch stimulus (or circle versus triangle cue stimulus in the case of the choose-match task), mean performance across trials relative to their distance (C) from the reversal between the presentation of match and nonmatch stimulus (or circle versus triangle cue stimulus in the case of the choose-match task) for novice versus expert training, the mean DIP (D) of each session over the course of training, and the weight index of five possible strategies (E) over the course of training, across all monkeys/tasks. All conventions are the same as Fig. 1.

**Supplementary Figure S3:**
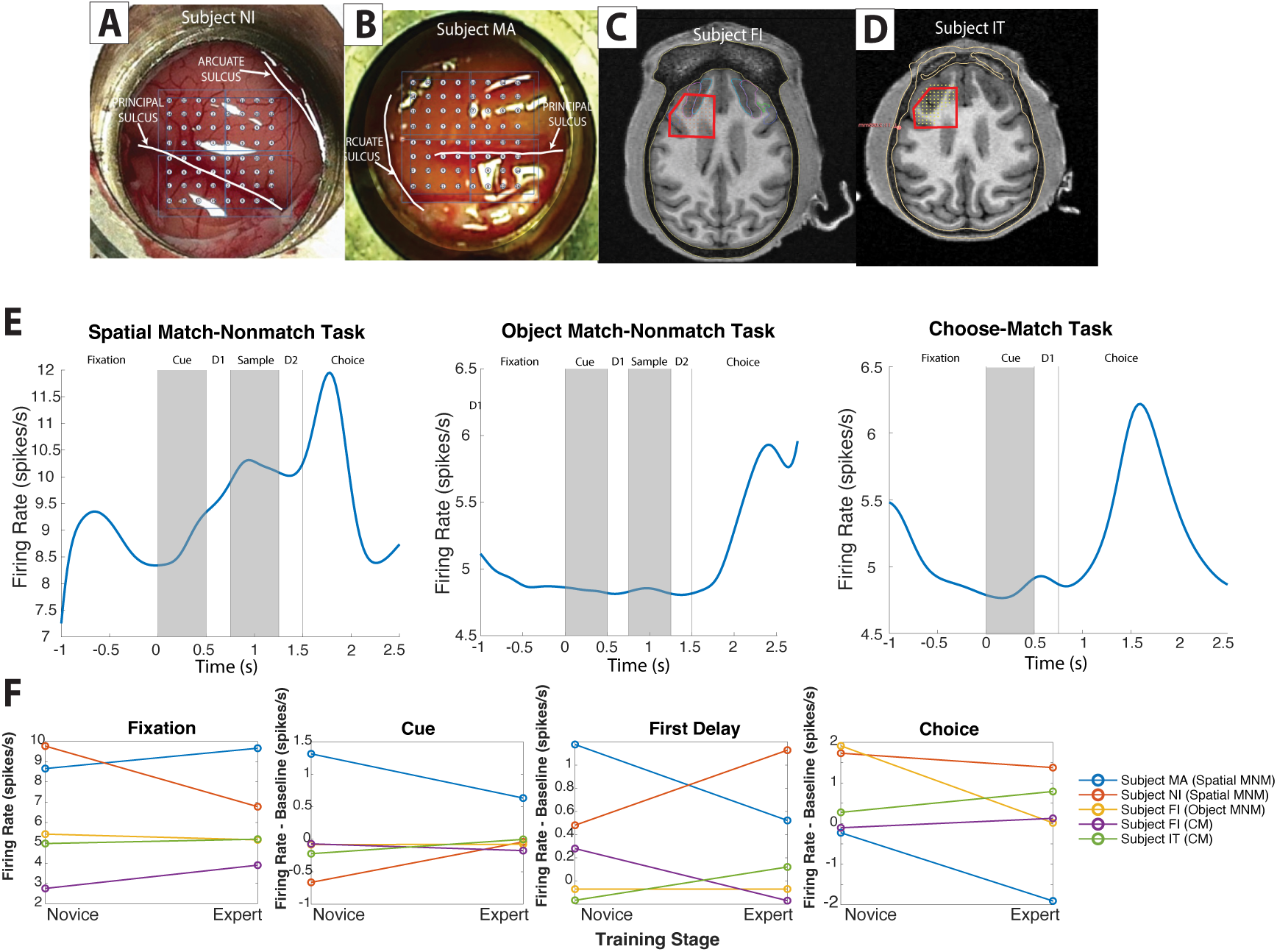
Neural recordings and activity. A. Position of the electrode array that was implanted into the dorsolateral and ventrolateral prefrontal cortex of Subject FI indicated in red. This was centered across the dorso-ventral and anterior-posterior axis—as defined by the line connecting the genu of the arcuate sulcus to the frontal pole. B. Position of the electrode array that was implanted into the dorsolateral prefrontal cortex of Subject MA, as indicated relative to the principal sulcus and arcuate sulcus. C. Position of the electrode array that was implanted into the dorsolateral and ventrolateral prefrontal cortex of Subject IT indicated in red. This was centered across the dorso-ventral and anterior-posterior axis—as defined by the line connecting the genu of the arcuate sulcus to the frontal pole. D. Position of the electrode array in the dorsolateral prefrontal cortex of Subject NI, as indicated relative to the principal sulcus and arcuate sulcus. E. Peri-stimulus time histogram (PSTH) plots depicting mean firing rate (y-axis) over the time course of the trial (x-axis) across all cells from each of our three experimental tasks. Gray bars indicate stimulus presentations. Time 0 represents the cue presentation. F. Mean baseline fixation and firing rates (y-axis) for after subtracting the mean baseline fixation activity for the cue epoch, first delay epoch, and choice epoch of each monkey/task (indicated by colors), across novice vs expert training (x-axis).

**Supplementary Figure S4:**
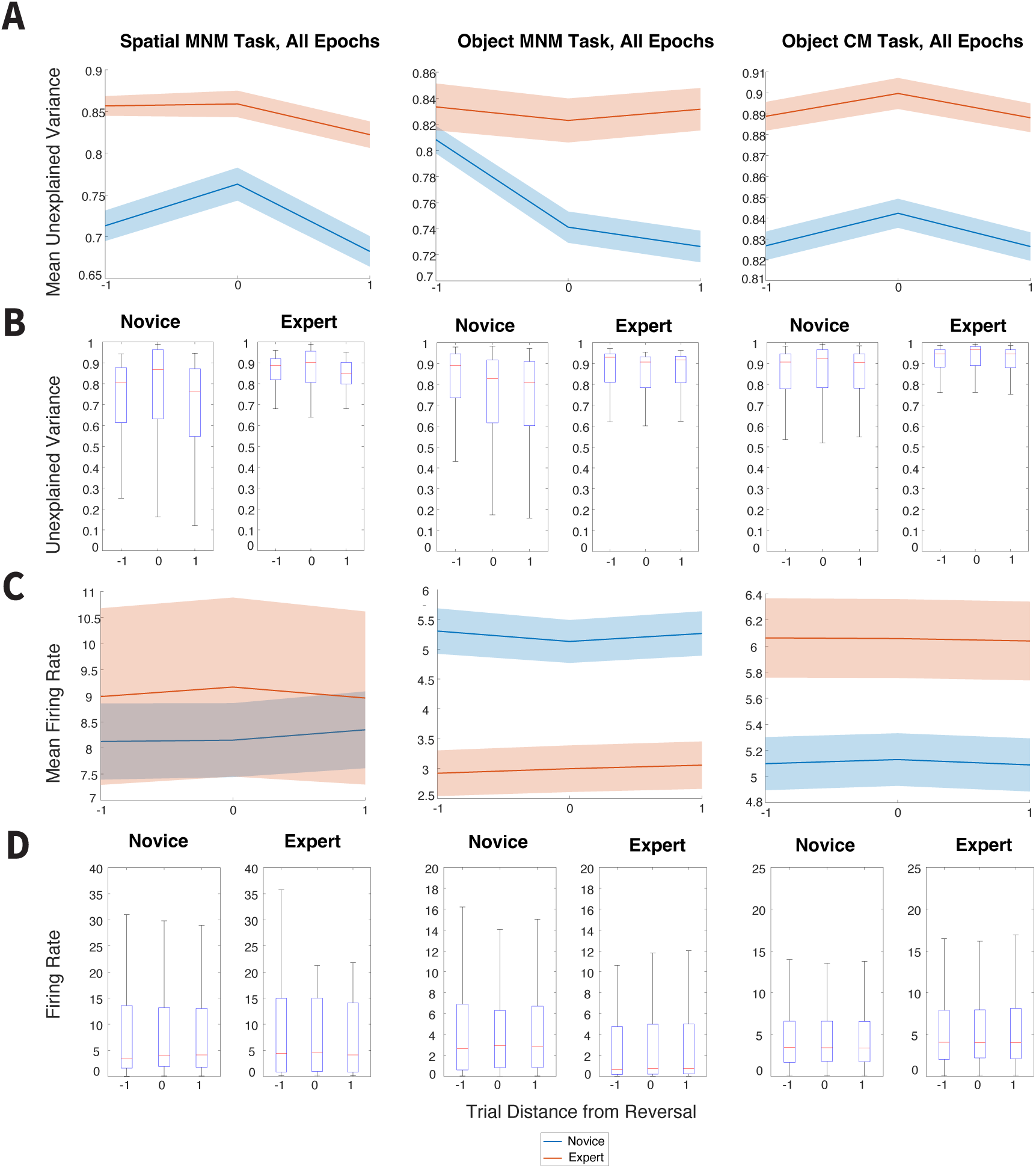
Expanded Unexplained Variance. Unexplained variance of firing rate (A-B) and the firing rate (C-D) in the spatial match-nonmatch (left), object match-nonmatch (middle), and object choose-match (right) tasks averaged across all epochs. Conventions are the same as in Fig. 3.

**Supplementary Figure S5:**
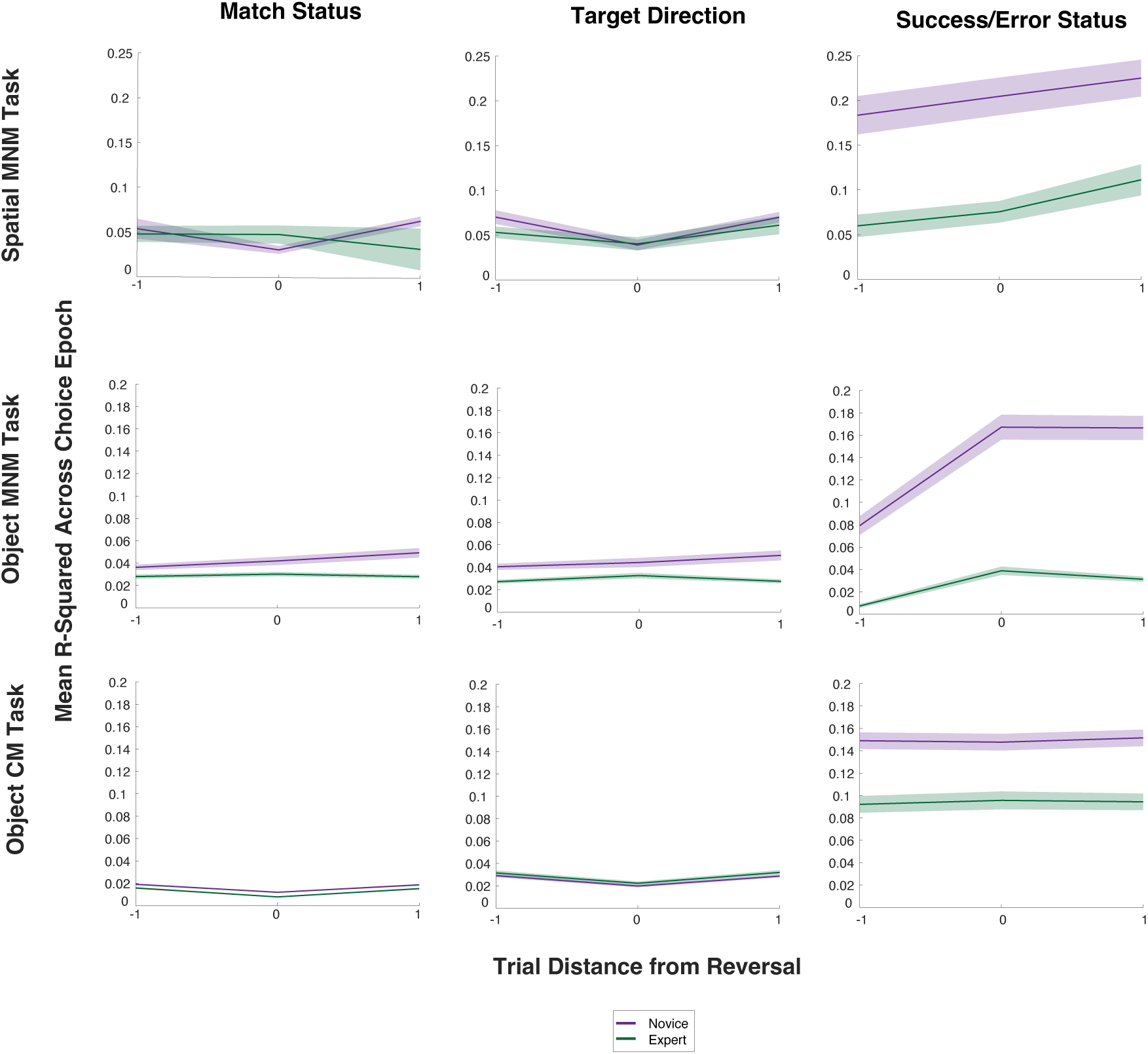
Partial correlation of task variables with firing rate. Mean partial R^2^ of firing rate from each task variable that was used in our regression model during the choice epoch across trials relative to their distance from the reversal. Shaded areas indicate standard error. Purple indicates novice while green indicates expert stage.

**Supplementary Figure S6:**
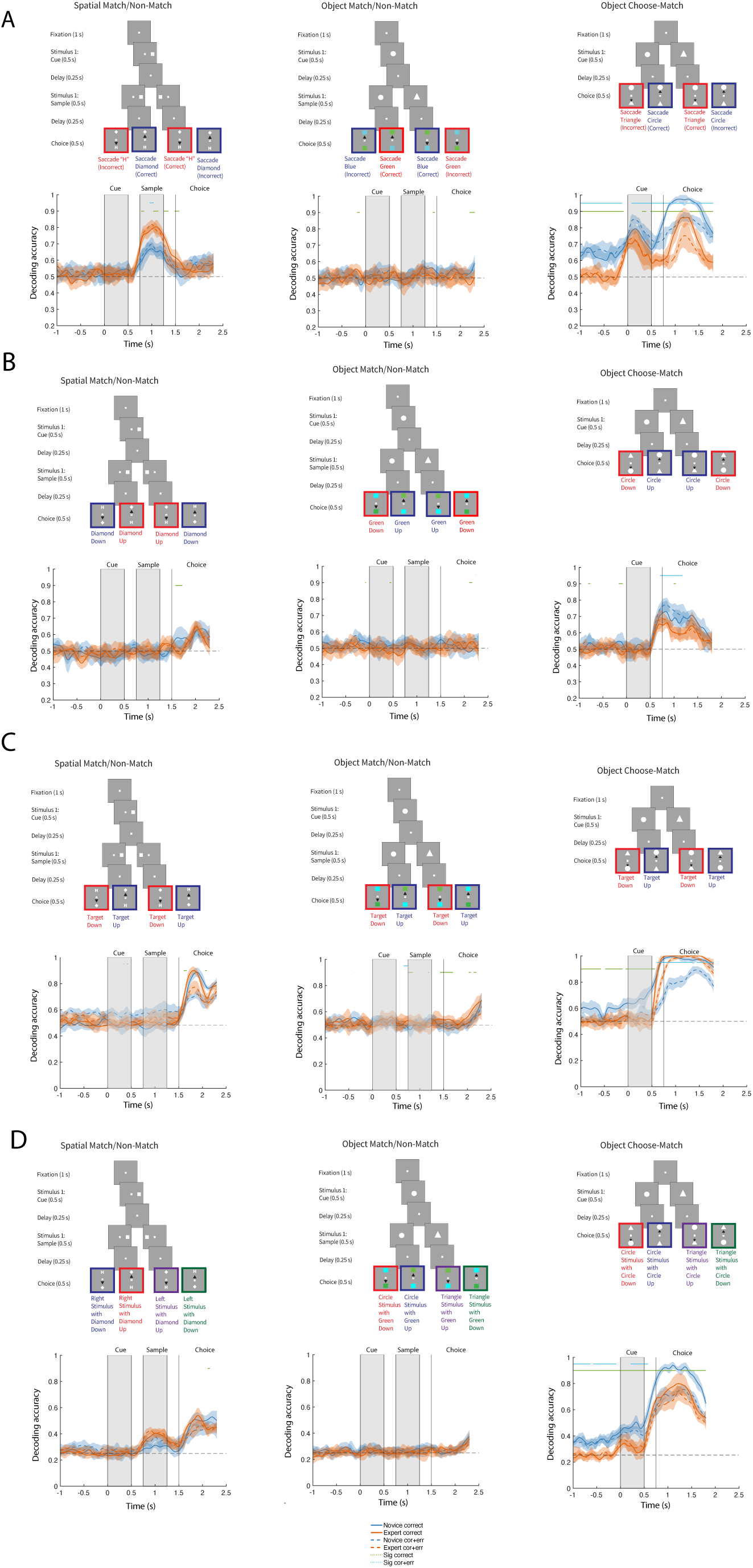
Expanded decoder analyses. Decoder performance for all task factors that were not included in Fig. 4. We performed our decoding analyses using an equal number of cells drawn from novice and expert training sessions. This was a total of 100 novice and expert cells from the spatial match-nonmatch task (left), 400 novice and expert cells from the object match-nonmatch task (middle), and 400 novice and expert cells from the object choose-match task (right). In each task, top diagrams illustrate the A. the monkey’s choice to saccade for a match regardless of whether this choice was correct, B. the configuration of the choice targets (i.e. whether a specific choice target appeared up or down regardless of whether this was the rewarded target), C. the direction of the rewarded target (regardless of whether the monkey chose to saccade to this target), and D. the combination of the presentation of match or nonmatch stimulus with the direction of the rewarded target. Bottom plots illustrate the accuracy of decoding (y-axis) this task factor from the novice (blue) versus expert (red) neurons over the time course (x-axis) of each task. Vertical black lines indicate each task epoch. Solid blue and red lines indicate the use of exclusively correct trials for training and testing data. Dashed blue and red lines indicate the use of both success and error trials for training and testing data. Shaded ribbons represent 95% confidence intervals. Significant differences between novice and expert decoding are marked overhead in green and cyan (permutation test, p < 0.01).

**Supplementary Figure S7:**
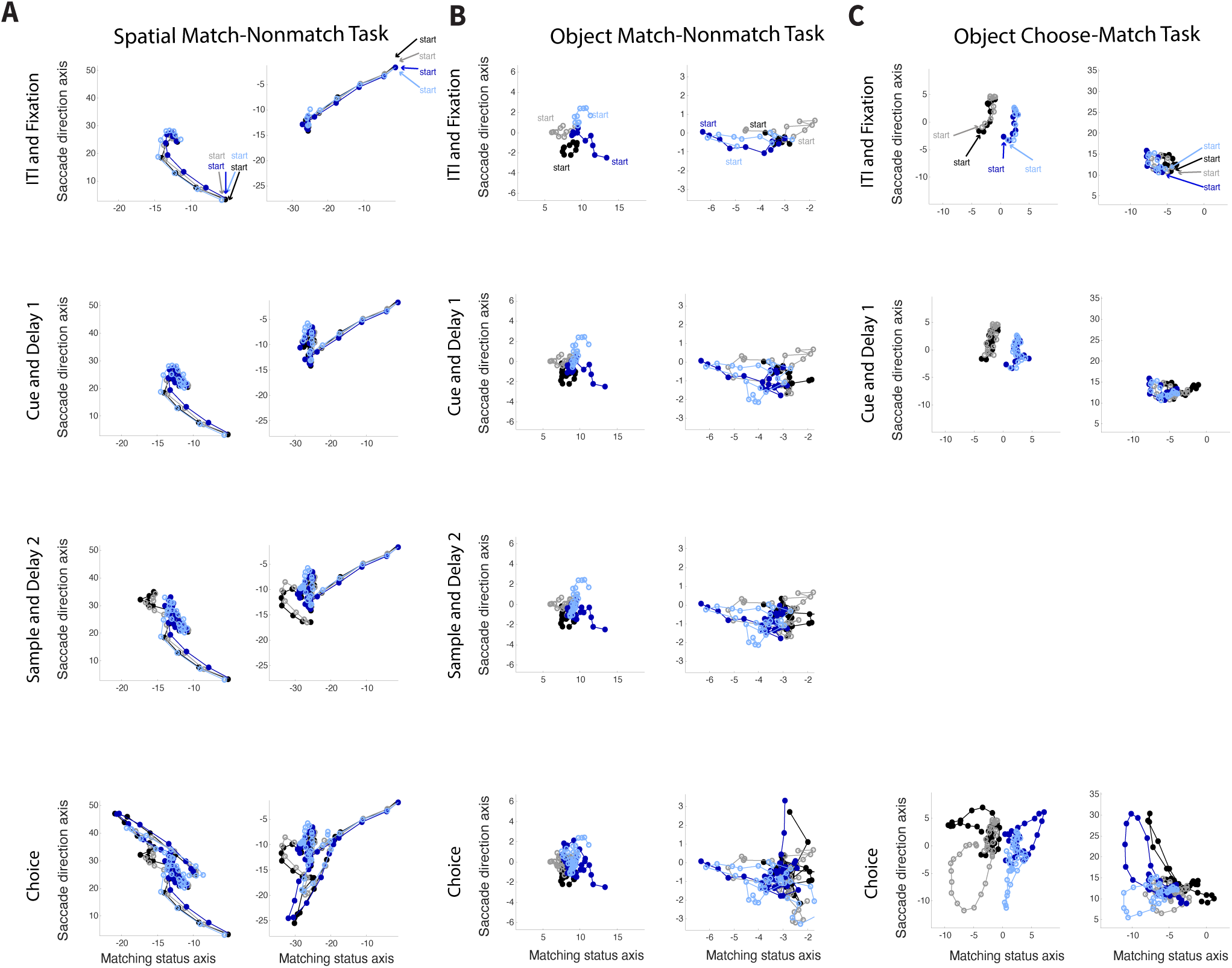
Expanded dynamics of populations responses in PFC. The average population response for a given condition and time is represented as a point in state space for A. the spatial match-nonmatch task, B. the object match-nonmatch task, and C. the object choose-match task. Responses from correct trials only are shown from 1000 ms before the start of the trial (inter-trial interval) to the end of the trial using a 400 ms sliding window, with a 100 ms step size (with overlap), and are projected into the two-dimensional subspace capturing the variance due to the combination of the presented stimulus (i.e. match or nonmatch in the spatial and object match-nonmatch tasks, and circle or triangle stimulus presentation in the object choose-match task) and the saccade direction of the rewarded choice target (up or down) task conditions for novice (left) and expert (right) training phases. Units are arbitrary. Each color indicates a different combination of the two pairs of possible task conditions. Starting points for each of the resulting neural trajectories is indicated by text and arrows in the same color. D. The distribution (y-axis) of the change in separability (x-axis) between empirical novice and expert neural trajectories (blue) versus a null distribution (orange) that was calculated from a pair of randomly shuffled surrogate datasets for the object choose-match task after aligning the mean position of the neural trajectories in the fixation period to account for the bias already present in the baseline. This revealed a clear increase in separability that mirrored the results of the spatial and object match-nonmatch tasks. Conventions are the same as Fig. 5e.

